# A genetic pathway encoding double-stranded RNA transporters and interactors regulates growth and plasticity in *Caenorhabditis elegans*

**DOI:** 10.1101/694414

**Authors:** Fabian Braukmann, David Jordan, Eric Alexander Miska

## Abstract

The environment and genes shape the development, physiology and behaviour of organisms. Many animal species can take-up double-stranded RNA (dsRNA) from the environment. Environmental dsRNA changes gene expression through RNA interference (RNAi). While environmental RNAi is used as a laboratory tool, e.g. in nematodes, planaria and insects, its biological role remains enigmatic. Here we characterise the environmental dsRNA receptor SID-2 to understand the biological function of dsRNA uptake in *Caenorhabditis elegans*. First we determine that SID-2 localises to the apical membrane and the trans-Golgi-network (TGN) in the intestine, implicating the TGN as a central cellular compartment for environmental dsRNA uptake. We demonstrate that SID-2 is irrelevant for nucleotide uptake from the environment as a nutritional (nitrogen) source. Instead RNA profiling and high-resolution live imaging revealed a new biological function for *sid-2* in growth and phenotypic plasticity. Surprisingly, lack of the ability to uptake environmental RNA reduces plasticity of gene expression. Furthermore, using genetic analyses we show that the dsRNA pathway genes *sid-2, sid-1* and *rde-4* together regulate growth. This work suggest that environmental RNA affects morphology and plasticity through gene regulation.

## Introduction

Gathering information about the environment is fundamental for the survival of an individual in an ecosystem. Recent studies show that one class of molecules that provides such information is RNA [1, 2]. Specifically, double-stranded RNA (dsRNA) is able to act in environmental communication by directly modulating gene expression in *Caenorhabditis elegans* [3, 4]. This process is known as environmental RNAi and further research identified that a number of animal species and fungi are able to do the same [5–10]. More recently, small RNAs were also shown to move between hosts and pathogens as part of an immunity arms race [11–13]. Nevertheless, the biological relevance and extend of RNA communication between organism remains poorly understood.

Environmental RNAi is an versatile and successful tool. Experimentalists use it in animals to investigate the function of genes by simply feeding them with bacteria overexpressing dsRNA with the sequence of the gene of interest [14]. Analogous, the method is successfully used for agricultural pest control, where genetically engineered plants express dsRNA with the sequence of an essential genes of the targeted parasite [15]. Delivery of dsRNA via a genetic modified organism (GMO) is efficient, but applications of environmental RNAi are not limited to GMO organisms. The development of non-GMO strategies, like artificial dsRNA nanoparticle sprays as topical antiviral for managing crop health [16] and large scale field application using artificial dsRNA sugar solution as antiviral tool in bee health [17], lay ground for cheaper and accelerated development of artificial dsRNA based applications and potentially an increased release of artificial dsRNA into the environment.

How such artificial dsRNA affects the ecology of targeted and non-targeted organism is an ongoing debate [18–22]. One of the great challenges is the identification of plausible pathway to harm, and additional basic research will continue to guide this process [23]. For example, field experiments helped to identify a spontaneous emergence of resistant to environmental RNAi in the target organism most likely due to mutation of genes in the dsRNA uptake pathway [24]. However, the effects of these mutations on development, physiology and behaviour of targeted and non-targeted organism remain uncharacterised.

Mutations blocking environmental RNAi have been identified in forward genetic screens in the model organism *C. elegans* and have led to a mechanistic understanding of artificial dsRNA uptake, transport and processing [25]. One such mutant defined an intestinal dsRNA receptor called <u>S</u>ystemic RNA Interference <u>D</u>efective-2 (SID-2) is essential for the uptake of artificial dsRNA [26]. Biochemical studies showed it is able to mediate the uptake of dsRNA and single stranded RNA forming only partial dsRNA structures in form of hairpins, but not the chemically similar DNA [27]. In addition, the entry of dsRNA is mediated by the endocytosis pathway as indicated by fluorescently labelled dsRNA soaking experiment in *Drosophila* tissue culture cells [28]. After the endocytosis of the dsRNA, a dsRNA transporter called SID-1 releases the dsRNA into the cytoplasm [29–31], where the dsRNA binding protein RDE-4 binds it [32]. RDE-4 together with the RNase III Dicer processes the dsRNA into small RNA duplexes, of which one strand is loaded into an Argonaute protein [33, 34]. The Argonaute small RNA complex interacts with complementary RNA sequences and mediates translational regulation and target RNA degradation [35–41]. In addition, they trigger the production of secondary small RNAs by RNA-dependent RNA polymerases (RdRPs), which amplify the gene-regulatory function [42–44]. Furthermore, the gene-regulatory effects can be inherited and last for several generations [45–48]. By this mechanism, artificial dsRNA can regulate gene expression and shape the phenotypic outcome over multiple generations.

Naturally occurring dsRNA has the potential to use the environmental RNAi pathway to alter phenotype. Within an organism, natural dsRNA can be synthesised during viral replication [49] and dsRNA with high sequence diversity is generated in many organisms, including *C. elegans* [50] and *E. Coli* [51], from their own genome most likely via anti-sense transcription [52–55]. Sequencing methods detected dsRNA with rich sequence diversity in microbial communities and water samples [56–60] indicating that dsRNA is a common part of an ecological system. Furthermore, natural dsRNA is found as well outside the organism. Honey bees feed their larvae with royal jelly, a complex milk like nutrient, rich in natural dsRNA sequences [61]. In addition, this feeding solution contains a protein with the ability to protect dsRNA from degradation [62] demonstrating a potential mechanism to preserve information encoded in dsRNA and possibly allowing communication via environmental RNAi. However, natural examples of dsRNA transport between organism and if they shape phenotype have not yet been observed yet. Furthermore, the biological roles of dsRNA uptake and transport mechanism present in many organism remain unclear.

In this study, we use genetic analysis of dsRNA uptake mutants *sid-2* to gain important insights into the biological function dsRNA uptake. We find that the SID-2 does not function in dsRNA uptake for nutritional purposes. However, SID-2 regulates endogenous small RNA pathways, animal growth and phenotypic plasticity in *C. elegans*, but not in *C. briggsae*. The regulation of *C. elegans’* growth also requires the dsRNA transporter SID-1 and dsRNA-binding protein RDE-4, which are known to act downstream of SID-2 in RNAi. Here we present the first evidene for a role of the dsRNA uptake pathway for morphology and phenotypic plasticity in *C. elegans*. Therefore this work conributes to a better understanding of environmental dsRNA uptake as a mechanism of communication between animals and the environment.

## Results

### *sid-2* is a conserved gene in the genus *Caenorhabditis*

The uptake of the environmental dsRNA by SID-2 is the first step for environmental RNAi in *C. elegans*. However, some closely related nematodes have lost the ability to take up dsRNA [26, 63]. For example, in the sister species *C. briggsae*, a homologue of *sid-2* is present, but unable to function in dsRNA uptake for environmental RNAi [26, 27, 63]. Newly available genome and transcriptome data allows us to study the evolution of *sid-2* in 25 *Caenorhabditis* species [64, 65]. *sid-2* is present in all but one analysed *Caenorhabditis* genomes (S1 Table) indicating that *sid-2* is a conserved gene in the *Caenorhabditis* genus. A phylogenetic analysis using the SID-2 protein sequences did not separate susceptible strains from those resistant to RNAi by feeding resistant strains (Fig 1A). Rather, the SID-2 sequences clustered into the known *Caenorhabditis* supergroups *drosophilae* and *elegans*, as well as into the groups *elegans* and *japonica*. Further, an alignment analysis of all SID-2 protein sequence indicate that the extracellular domain is less conserved than the transmembrane domain and intracellular domain (S1A Fig) indicating that functional difference are more likely due to changes in the extracellular domain. This result indicates no common amino acid change in SID-2 can explain the susceptibility or resistant to environmental RNAi.

**Fig 1.**
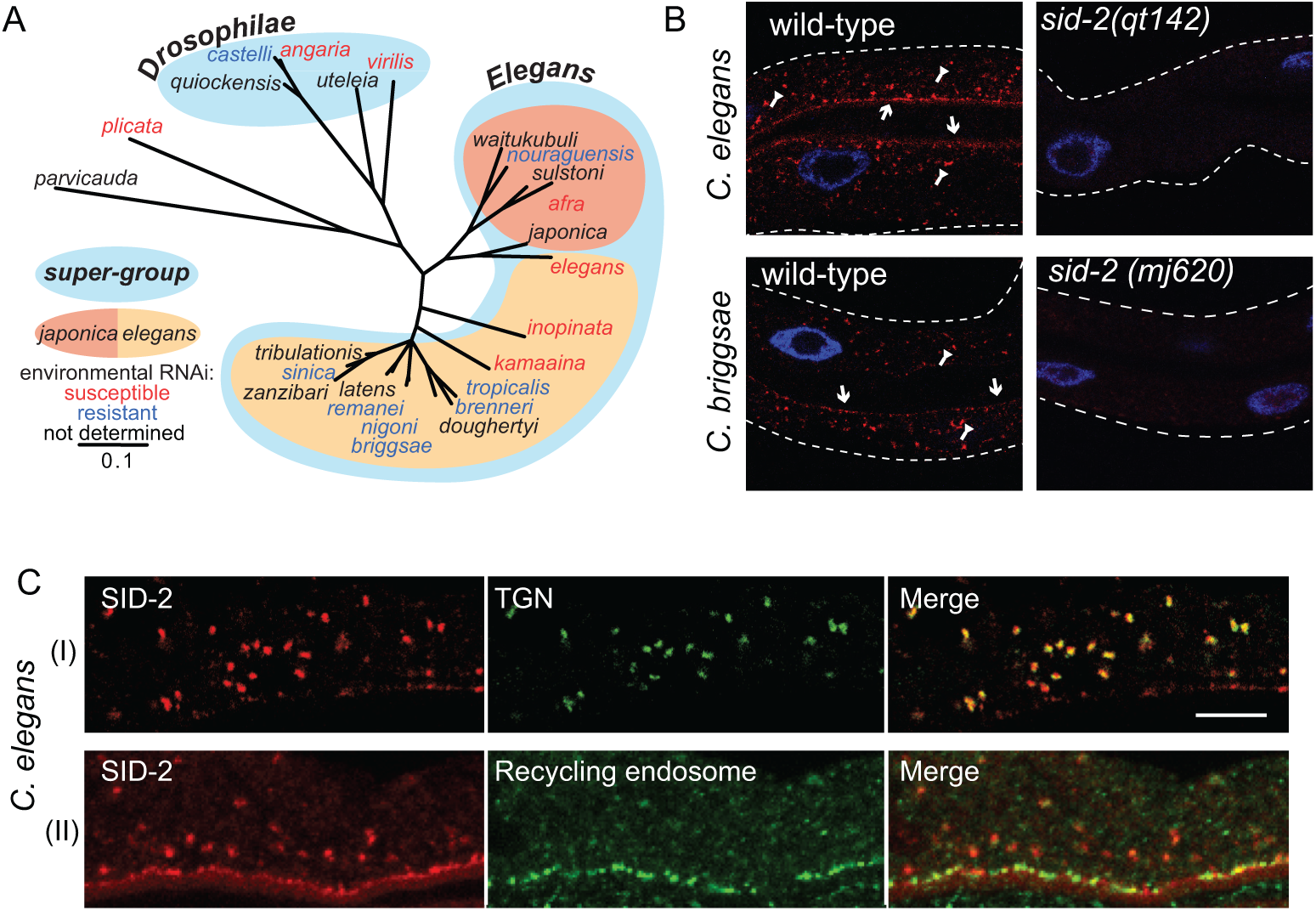
SID-2 has a conserved localisation at the intestinal apical membrane and at the TGN *in vivo*. A) Maximum likelihood tree (100 bootstraps) using the most conserved region of the potential dsRNA receptor SID-2 of 25 *Caenorhabditis* nematodes. Environmental RNAi susceptibility (red), environmental RNAi resistant (blue). B) Confocal microscopy image of *C. elegans* and *C. briggsae* dissected adult intestines after SID-2 immunofluorescence staining. DNA was stained using DAPI (blue). In *C. elegans* and *C. briggsae* wild-type, SID-2 was detected close to the lumen (pointed arrows) and in punctuated foci (flat arrows), but not in *sid-2* mutants. Doted line outlines intestine. Representative image is shown (n = 3). C) Confocal microscopy image of *C. elegans* dissected adult intestines expressing flourescent protein fusions to SYN-16 and RAB-11 after SID-2 and GFP co-immunofluorescence staining. SID-2 antibody staining in red (left column), GFP antibody staining in green (middle column) and merged image (right colum). (I) TGN (YFP::SYN-16) and (II) recycling endosomes (GFP::RAB-11). Representative image is shown (n = 3), scalebar = 5 *µ*m.

### SID-2 localises in the adult intestine close to the apical membrane and the trans-Golgi network in the genus *Caenorhabditis*

Because, previous reports showed that *C. briggsae* SID-2 does not function in artificial dsRNA uptake, we wanted to understand if the subcellular localisation of SID-2 is responsible for the deficiency in dsRNA uptake. To address this question, the subcellular localisation of SID-2 was analysed using SID-2 immunofluorescence staining in dissected intestines of *C. elegans* and *C. briggsae* wild-type and *sid-2* mutant animals (S1B, C Fig). First, we created *C. briggsae’s sid-2* mutant by CrisprCas9 genome editing (S1C Fig, S2 Data) and then performed the immunofluorescence staining with a custom made SID-2 antibody, predicted to target the intracellular domain of SID-2 (S1B Fig). In both species, SID-2 localised close to the apical membrane, in line with previous reports [26], in addition it localised to numerous foci in the cytoplasm (Fig 1B). SID-2 was not detected in *C. elegans* nor in *C. briggase sid-2* mutant animals (Fig 1B, S1D Fig). In strains overexpressing subcellular compartment markers fused to a fluorescent protein, SID-2 and fluorescent protein co-immunofluorescence staining revealed that SID-2 co-localises with the recycling endosomes marker RAB-11 close to the apical membrane, with the trans-Golgi-network marker SYN-16 (Fig 1C) [66] and adjacent to the middle stack Golgi marker MANS, but not with the autophagosomes marker(LGG-1) nor the late endosome marker RAB-7 (S1E Fig). This is the first time endogenous SID-2 localisation has been analysed. The novel localisation of SID-2 at the TGN indicates the importance of the TGN for the function of SID-2 and the localisation of SID-2 at the apical membrane suggests that SID-2 is in contact with the environment at the intestinal apical membrane in vivo.

### *sid-2* does not function in dsRNA uptake for nutritional reasons

Because SID-2 localisation is conserved in *C. elegans* and *C. briggsae*, but mediates artificial dsRNA uptake only *C. elegans*, we wondered if SID-2 conserved function is enhancing the uptake of dsRNA for nutritional purposes. Since the nutrient rich laboratory environments might mask such a function, we use a *C. elegans* strain with a compromised PYRimidine biosynthesis pathway *pyr-1* [67] to test if SID-2 takes up dsRNA for nutritional reasons. Specifically, we asked if exogenous sources of pyrimidines contribute nutritionally in the *pyr-1* sensitised background, and if the uptake of exogenous sources requires *sid-2* (Fig 2A). In this experiment, *C. elegans* wild-type, *sid-2*, *pyr-1* and *pyr-1;sid-2* double mutant animals were grown on *E. coli* bacteria in four conditions supplemented with exogenous pyrimidine in different ways (None, Uracil, long dsRNA and short dsRNA) (Fig 2B, S2A Fig). First, we determined the hatching rate under the control condition, when no pyrimidine supplement was provided, 100 % of the wild-type and *sid-2* embryos hatched, indicating *sid-2* does not affect embryonic development on its own and that sufficient pyrimidines are present (Fig 2B). As previously demonstrated, *pyr-1* mutants showed a severe embryonic development defect with a hatching rate of *≈* 10% [67], similar to *pyr-1; sid-2* double mutants (Fig 2B) indicating *sid-2* does not affect embryonic development in a *pyr-1* mutant. Next, the addition of exogenous Uracil rescued the embryonic development defect in *pyr-1* mutant animals, indicating that environmental pyrimidine can contribute nutritional value. In *sid-2, pyr-1* double mutants the embryonic hatching rate was similarly rescued, indicating that *sid-2* is not required for the uptake of Uracil. (Fig 2B). Finally, we overexpressed long dsRNA and short dsRNA in the *E. coli* bacteria (Fig 2B, S2A Fig). In these condition long and short dsRNA was equally or more abundant then ribosomal RNA (S2B Fig). Embryonic development was benchmarked in wild-type animals and did not differ in *sid-2* mutant animals. In *pyr-1* mutants, the expression of long or short dsRNA was able to improve the embryonic development to a hatching rate of 60% and 70%, respectively (Fig 2B,S2A Fig) and similar hatching rate were observed in *pyr-1;sid-2* double mutants. Thus, the environmental dsRNA contributed to nutrition independently of *sid-2*.

**Fig 2.**
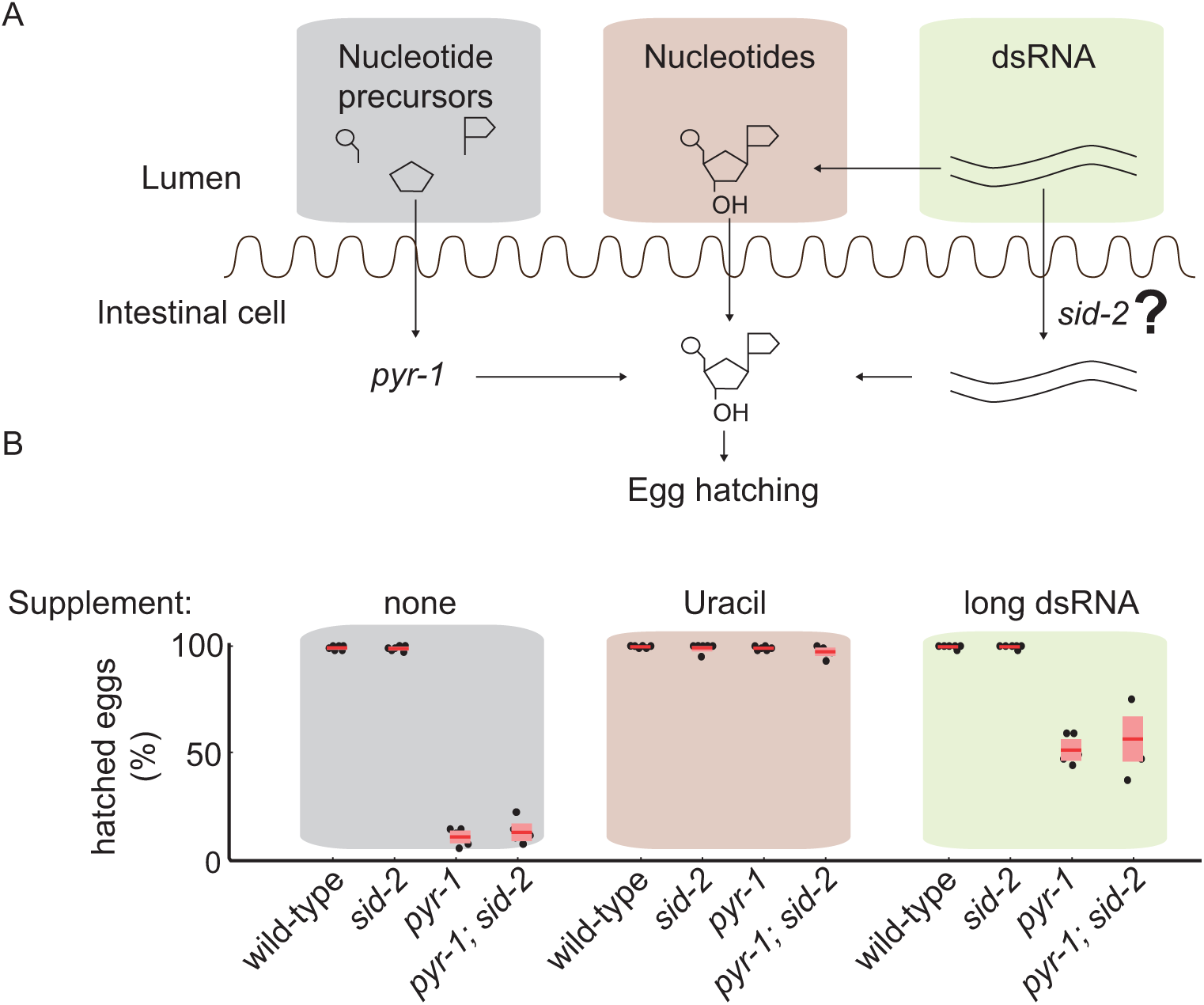
The conserved dsRNA receptor SID-2 does not enhance dsRNA uptake for nutritional reasons in *C. elegans*. A) Model of nucleotide uptake and synthesis for egg hatching. Egg hatching depends on sufficient pyrimidines within the animal. Pyrimidines can be synthesised by *pyr-1*, taken up from the environment or potentially derived from environmental dsRNA taken up by SID-2. B) The effect of pyrimidine supplement and dsRNA expression on egg hatching in wild-type, *sid-2(qt142)*, *pyr-1(cu8)* or *pyr-1(cu8);sid-2(qt142)*. Black dimonds represent percent hatching of eggs of individual experiments (n = 3 with 2 technical replicates). Red lines indicate the mean. Red boxes indicate the 95% confidence interval of the mean.

The conclusion of the above assay relies on the ability of *pyr-1* mutants to take up dsRNA. To show that *pyr-1* mutants can take up dsRNA, we performed RNAi by feeding since that relies on dsRNA uptake. Specifically, RNAi was fed in an environment supplied with exogenous Uracil. On negative control vector, all genotypes showed 100 % hatching efficiency indicating that in this condition embryonic development is unaffected (S2C Fig). Next, we fed RNAi against POsterior Segregation-1 (*pos-1*), a gene essential for embryo development, again in the presence of exogenous Uracil. The drastically reduced hatching (*≈* 10%) of wild-type and *pyr-1* mutant animals hatched, showing that animals are capable to take up dsRNA and mount an RNAi response. Finally, all strains with a *sid-2* mutation had a hatching rate of 100% indicating that they were unable to take up dsRNA (S2C Fig). Therefore, we concluded that *pyr-1* mutants can take up dsRNA, and *sid-2* does not enhance dsRNA uptake for nutritional purposes.

### *sid-2* reduces growth rate but increases phenotypic plasticity

Because artificial environmental dsRNA can induce gene regulation, naturally occurring dsRNA might induce directly or indirectly induce gene expression changes, which we can measure. In an attempt to detect such changes, we performed transcriptome analysis in embryos of wild-type animals and two *sid-2* mutants deficient in dsRNA uptake (S1 Fig, S2 Table) fed on standard lab bacteria. First, we confirmed using differential expression analysis that the individual *sid-2* mutants were similar to each other (S3A Fig). Next, we measure transcriptome differences between each *sid-2* mutant and the wild-type embryos (S3B and C Fig). In *sid-2(qt142)* mutants, 407 transcripts had expression different from wild type (FDR *<*0.01) (S3D Fig). In *sid-2(mj465)*, a slightly higher number of transcripts (569) had a different expression than in wild type (FDR *<*0.01) (S3D Fig). A total of 314 common different transcripts were detected (S3D Fig), this significant overlap (Hypergeometric Test *≈* 0) argues that the *sid-2* mutants behave similarly, therefore we combined the individual *sid-2* mutant transcriptomes to gain additional statistical power in differential expression analysis against the wild-type embryo transcriptomes. In this analysis, we identified 935 significantly differentially expressed genes (FDR *<*0.01) in *sid-2* mutant embryo transcriptome (Fig 3A). In a summarising gene ontology enrichment analysis [68], a significant overlap was found with genes functions like structural constituents of the cuticle, collagen trimmer and cytoskeleton (S3E Fig). These three biological processes are associated with the *C. elegans* exoskeleton which undergoes drastic rearrangements during development suggesting that SID-2 may alter embryonic development.

**Fig 3.**
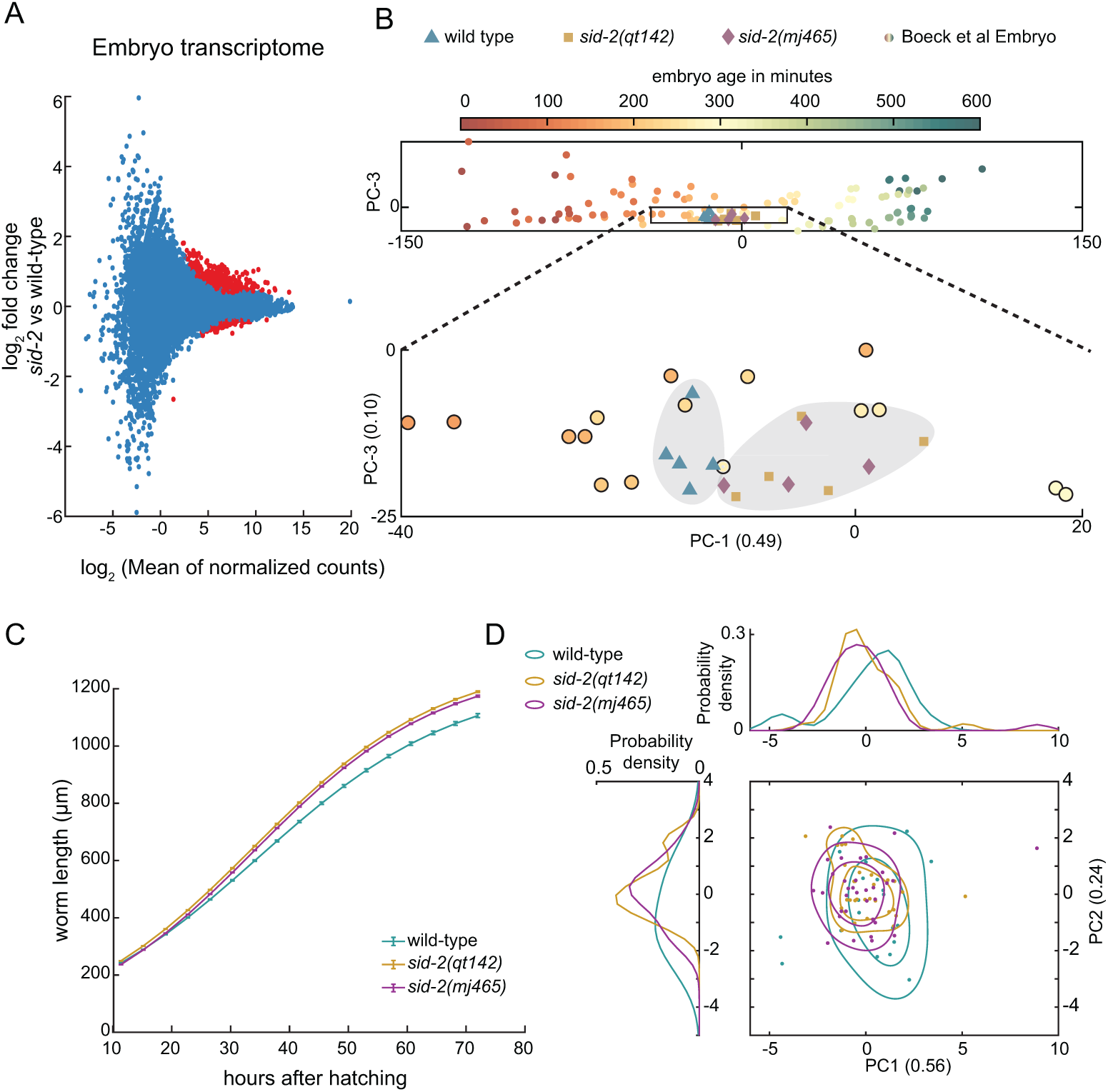
*sid-2* slows growth rate and increases phenotypic plasticity. A) MA plot visualising embryo transcriptome comparison of wild type (n = 5) and *sid-2* mutants (total n = 9, *sid-2(qt142)* n = 4, *sid-2(mj465)* n = 5). Each red circle represents a statistically significant (DE) transcript (FDR *<*0.01). B) Embedded wild type and *sid-2* mutant embryo transcriptome in principle component space of embryo development data (Boeck et al 2016). C) Growth curve visualising body length from hatching to egg laying adult of wild-type animals (n = 20) and *sid-2(qt142)* (n = 31), *sid-2(mj465)* (n = 33) mutants, line represent median and error-bar represent the 95% confidence interval of the median. D) Principle component analysis representing the combined phenotypic data along the first and second principle component of the analysed wild-type animals (n = 20) and *sid-2(qt142)* (n = 31), *sid-2(mj465)* (n = 33) mutants. Individual circles represent aggregated phenotypical data of a individual animal. The line represent the 33% and 66% contour line. Probability density estimate of the phenotypic data is plotted left and top of the PCA plot.

To investigate if *sid-2* mutant embryos develop faster or slower than wild-type embryos, we embedded our transcriptome data in the lower dimensional space generated from a time series of embryo transcriptomes [69]. The principle component analysis ordered early to late embryonic transcriptomes from the left to the right along the first principle component (Fig 3B top, S3F Fig). The wild-type and *sid-2* transcriptomes were projected in this lower dimensional space. The wild-type transcriptomes clustered further left, whereas the *sid-2* mutant transcriptomes clustered further to the right (Fig 3B) indicating that *sid-2* mutant embryos are more similar to older embryos, and therefore advanced in development compared to wild-type embryos.

This analysis led us to investigate whether *sid-2* affects growth rate at other developmental stages. Therefore, we measured the body length using single animal imaging of wild-type and *sid-2* mutant animals throughout the development from hatched larvae to adulthood. The growth rate, estimated by a logistic function (S3 Table), was slower in wild-type animals than in *sid-2* mutants (Fig 3C), suggesting that the present of SID-2 slows development perhaps do to dsRNA uptake.

From the same individuales the ex-utero development time and the time from hatching until the first egg was laid were collected (S3 Table). Together with the three estimated parameters from the logistic function (logistic max, logistic rate, logistic shift), we constructed a phenotypic space. To explore this space, we displayed the phenotypic space in two dimensions using all three combinations of the first, second and third principle components (Fig 3D, S4A and B Fig). We observed that the data points of wild-type animals had a wider spread compared to *sid-2* mutant animals (Fig 3D) indicating that wild-type animals are more variant that *sid-2* mutant animals (S3 Table). To test if the phenotypic plasticity of *sid-2* mutants was different from wild-type animals, we performed a statistical analysis testing for equality of the variation of the projection phenotypic data on the first four principle components containing more than 99% if the variance comparing *sid-2* mutant and wild-type animals (Box’s M tests p *<*0.0016). This results suggest that SID-2 increases phenotypic plasticity.

### SID-2 is expressed in all larvae stages and adults

To understand at what stages of development SID-2 may affect growth and phenotypic plasticity, western blot analysis was performed throughout development. The specificity of the SID-2 antibody was tested by comparing wild-type adults and *sid-2(qt142)* null mutant adults (S5A Fig). SID-2’s predicted molecular weight is 33 kDa, however bands were detected at *≈* 40 kDa and at *≈* 20 kDa that were absent in *sid-2* mutants. At *≈* 40 kDa two bands were detected indicating that two isoforms of SID-2 exist, potentially with different post-translational modifications (e.g. glycosylation, a modification placed in the Golgi [70]). The smallest band at *≈* 20 kDA is potentially a degradation product of full length SID-2. The band observed at *≈* 60 kDa was detected as well in *sid-2* mutants, and therefore likely result from non-specific binding to the SID-2 antibody. SID-2 expression was detected in all tested developmental stages (S5B Fig) suggesting that SID-2 functions throughout the animals life.

### *sid-2* mutants have abnormal endogenous small RNA populations

Next we assessed the effect of *sid-2* mutation on endogenous small RNA populations. We performed RNA sequencing of primary and secondary siRNAs of wild-type and *sid-2* L4 animals. Loss of SID-2 has a significant effect on both populations (S6A Fig). We hypothesise that this is due to competition with environmental RNA in the wild type. To identify a potential exogenous RNA responsible for the *sid-2* phenotype, we sequenced long and short RNA of wild-type and *sid-2* L4 animals. We then compared the amount of *E. coli* derived reads from wild-type and *sid-2* mutant animals. No significant difference in the total amount of bacteria-derived RNA was detected (S6B Fig). Next, we asked if *sid-2* is required for the uptake of a specific transcript. Using differential expression analysis we were unable to identify *E. coli* transcripts that were significantly lower in abundance in *sid-2* mutants than in wild-type animals (S6C Fig). Together these experiments indicate that, we were unable to identify a potential causative environmental RNA, suggesting that body length could be regulated by dsRNA that is undetectable by the current sequencing method or alternatively by a mechanism unrelated to dsRNA.

### *C. elegans* growth is affected by a genetic pathway including the dsRNA interacting genes *sid-2*, *sid-1* and *rde-4*

To further investigate the involvement of dsRNA in the regulation of growth and phenotypic plasticity, we hypothesised that mutations in downstream dsRNA transport and dsRNA processing genes may cause similar morphological changes. Due to the low throughput of the life time image analysis platform, we decided to measure L1 length at hatching as a proxy for growth, allowing us to measure more individuals and more genotypes in less time. We tested if *sid-2* affected the morphology of freshly hatched larvae using a high resolution and magnification microscopy setup combined with image analysis to measure the length at birth. This analysis indicated that wild-type animals and known mutants with elongated body length *long-2* (*lon-2*) [71] have a mean body length at hatching of *≈* 179 *µ*m and *≈* 190 *µ*m, respectively (Fig 4A grey highlighted area). In contrast, the mean length at hatching of two strains carrying independent *sid-2* mutant alleles is *≈* 204 *µ*m, significantly longer. Restoration of *sid-2* function via overexpression in a *sid-2* mutant rescued body length comparable to wild-type animals (Fig 4A brown highlighted area). Together, these experiments suggest that *sid-2* affects body length.

**Fig 4.**
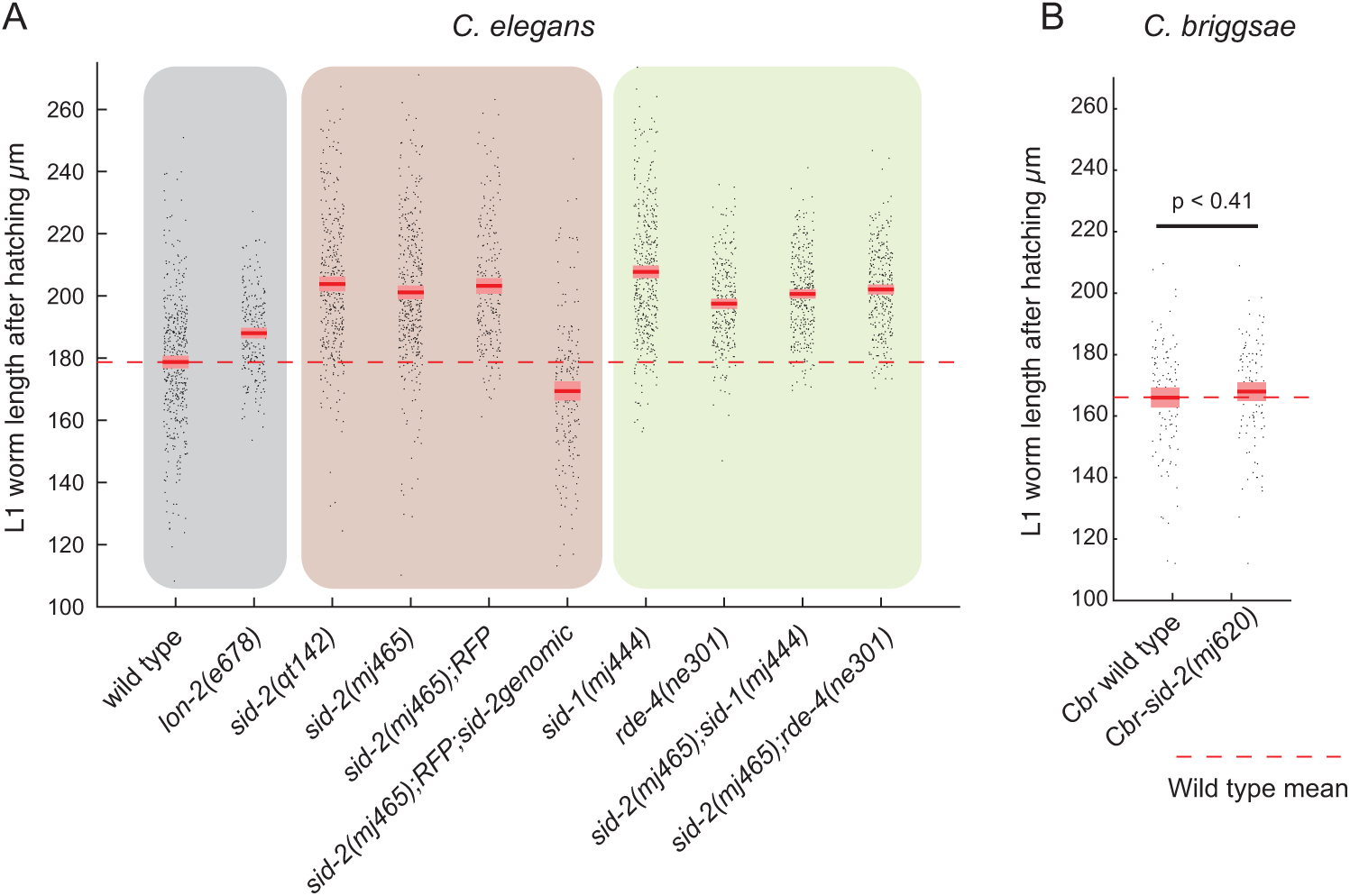
dsRNA interacting genes affect body length. A) Body length at hatching of *C. elegans* wild-type and indicated mutant animals. Black dots represent individual animals. Red lines indicate the mean, red boxes indicate the 95% confidence interval of the mean. Aggregate data from at least three experiments per genotype is shown. Individual experiments are shown in (S7 Fig). B) Body length at hatching of *C. briggsae* wild-type and *sid-2* mutant animals. (Two sample T-test p *<*0.41)

Next, we wanted to know if the changes in morphology are caused by dsRNA, therefore we performed the length measurement in additional mutants also deficient in dsRNA transport (*sid-1*) and dsRNA processing (RNAi DEfective-4 (*rde-4*)). For both *sid-1* and *rde-4* mutant animals, body length at hatching was significantly longer compared to wild-type animals and similar to *sid-2* mutant animals (Fig 4A green highlighted area). We performed an epistasis assay to determine if the effects on body length of *sid-2*, *sid-1* and *rde-4* mutants are due to the same genetic pathway and did not observe an additive effect (Fig 4A green highlighted area) suggesting that the three dsRNA interacting genes form one genetic pathway to alter length.

To further support the involvement for a dsRNA in the body length, we performed the length measurement in *C. briggsae* wild-type animals and compared them to *C. briggsae sid-2* mutants animals. The body length did not differ between wild-type and *C. briggsae sid-2* mutant animals (Fig 4B) showing that *sid-2* does not regulate body length in *C. briggsae* and suggesting that *sid-2* in *C. elegans* regulates body length via exogenous dsRNA that is yet to be identified. Overall, this experiment suggests that in *C. elegans sid-2*, *sid-1* and *rde-4* form a pathway which potentially processes exogenous dsRNA to regulate animal morphology (Fig 4C).

## Discussion

Artificial environmental dsRNA is extremely potent to induce gene regulation and phenotypic changes. In such artificial circumstances dsRNA is often used at extremely high concentrations, *e.g. ≈* 25% of the environmental RNA is double stranded (S2B Fig). However, weather animals take up naturally occurring RNA to regulate their phenotype is unknown. Here we study the intestinal dsRNA receptor SID-2 and show, that *sid-2* is a conserved gene in *Caenorhabditis*, which does not take up dsRNA for nutritional reasons. SID-2 is expressed throughout the larvae stages and in the adult animals and localises at the TGN and near the apical membrane. SID-2 regulates phenotypic plasticity, and together with the dsRNA transporter SID-1 and dsRNA binder RDE-4 body length. Here we discuss how growth and phenotypic plasticity can be regulated by *sid-2*, *sid-1* and *rde-4*, the potential sources of exogenous dsRNA, and the biological meaning of the observed phenotypic consequences.

### Unpredicted increase in phenotypic plasticity by SID-2

Mathematical modelling of gene networks and gene expression analysis of *E. coli* mutants reported that most, and perhaps all, mutation increase phenotypic plasticity when functionally compromised [72]. Further, experiments in the multicellular organisms *C. elegans* showed that mutations in individual components of a small gene-network lead as well to increase plasticity in gene expression and different phenotypic outcome [73]. Contrary to what theory and experimental data predict, we show that mutations in *sid-2* reduce surprisingly the phenotypic plasticity indicating that *sid-2* is one of the biological exception increasing phenotypic plasticity. Therefore, understanding the mechanism how *sid-2* reduces the phenotypic plasticity could uncover unique insides into biology.

### Mechanism of the regulation of growth and phenotypic plasticity

SID-2, SID-1 and RDE-4 act together to initiated gene silencing induced by exogenous artificial dsRNA. The exogenous artificial dsRNA is taken up by SID-2, transported into the cytoplasm by SID-1 and processed by RDE-4 and Dicer into small RNA [26, 27, 29, 32, 74, 75]. Here we use genetics to show that *sid-2*, *sid-1* and *rde-4* together regulate body length. Since *sid-2*, *sid-1* and *rde-4* have been shown to be involved in dsRNA processing [26, 27, 29, 32, 74, 75], we speculate that naturally occurring dsRNA is transported and processed by these proteins to regulate growth. Furthermore, due to the localisation of SID-2 in the apical intestine membrane we suspect, that dsRNA from the environment is regulating growth (Fig. 5).

**Fig 5.**
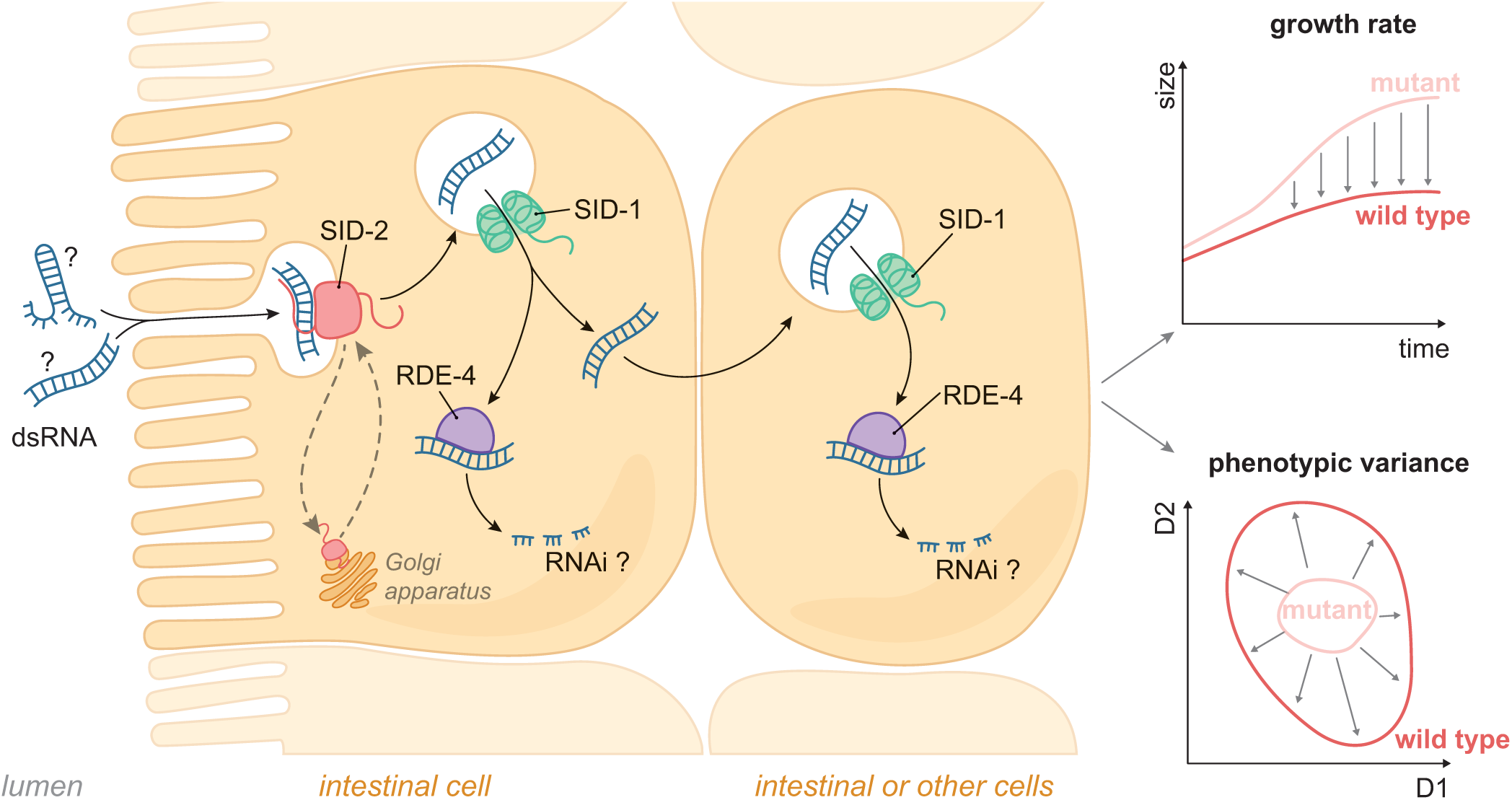
Working model of naturally occurring dsRNA regulating animal growth and phenotypic plasticity. Working model of exogenous dsRNA affecting animal growth via *sid-2*, *sid-1* and *rde-4*. Unidentified dsRNA is taken up by dsRNA receptor SID-2. dsRNA enters the cytoplasm via the dsRNA channel SID-1. There it interacts with the dsRNA binding protein RDE-4 entering the RNAi machinery. Growth and phenotypic robustness are affected by either targeting specific gene or unbalancing endogenous small RNAs. Furthermore, SID-2 might cycle between the TGN and the apical membrane.

### Origin of potentially exogenous dsRNA

Where does such dsRNA come from? Is it double stranded RNA from other *C. elegans* living in proximity, or is it dsRNA secreted from itself, for examples in form of extracellular vesicles [76, 77]. Perhaps, it is double stranded RNA from the bacteria that *C. elegans* are eating? Both are exciting models. The identification of such RNA is the next important step, followed by the validation that this RNA is causative for the observed phenotypic effects. If the RNA from *E. coli* would cause phenotypic consequences, it would be a stunning phenomenon. *E. coli* is no a natural food source for *C. elegans*, however if it is able to do induce the phenotype, than it would be likely that the RNA is common to many bacteria, suggesting environmental RNA mediated changes to the phenotype could be very common phenomena.

### Relevance of altered growth and phenotypic plasticity

The identified phenotypic consequences in laboratory conditions have limited predictive power to the real biology of free living *C. elegans*. Without an understanding of natural environmental conditions, the meaning of the reduced phenotypic plasticity remains unknown. Potentially, the effects of *sid-2* mutations are a small price for the ability to sample the environment for more beneficial dsRNA (e.g. one that confers immunity against a virus) [78]. Nevertheless, the rigours phenotypic analysis of dsRNA uptake mutant in highly controlled environments might help to design future studies guiding risk assessment of environmental RNAi applications. Overall, our results demonstrate for the first time the phenotypic consequences of the loss of the environmental RNAi pathway outside of experimental RNAi itself. Our work provides the first evidence of dsRNA-mediated control of organismal plasticity. Future investigation should focus on determining on how a more natural environment shapes *C. elegans* through RNA. We expect that such RNA-based interactions will be of broad relevance in many species.

## Methods

### Nematode culture and strains

*C. elegans* was grown under standard conditions at 20 °C. Bristol N2 was used as wild-type strain [71]. The *E. coli*strain HB101 [*supE44 hsdS20(rB-mB-) recA13 ara-14 proA2 lacY1 galK2 rpsL20 xyl-5 mtl-1*] was used as food source, except HT115[*F-, mcrA, mcrB, IN(rrnD-rrnE)1, rnc14::Tn10(DE3 lysogen: lavUV5 promoter -T7 polymerase*] was used in the pyrimidine supplement experiment and RNAi experiments. Both bacteria strains were obtained from the Caenorhabditis Genetics Center, University of Minnesota, Twin Cities, MN, USA.

### RNAi experiments

Empty vector, *pos-1* and *rpb-2* bacterial feeding clones were a kind gift from J. Ahringer’s laboratory. Bacteria were grown in LB Ampicillin for 6 hours, then seeded onto 50 mm NGM agar plates containing 1 mM IPTG and 25 µg/ml Carbenicillin. A volume of 200 µl bacterial culture per plate was used and left to dry for 48 hours. Further details on RNA interference are described in Kamath et al., 2003. L1 larva were synchronised by bleaching and transferred onto a RNAi plates and body length compared to wild type was assayed after 72 hours. For RNAi of *pyr-1* mutant strains, 0.5% Uracil was added to the NGM IPTG Carb plates.

### Tree construction

SID-2 protein sequences were obtained from www.caenorhabditis.org and www.wormbase.org as indicated in the supplemental file [64, 65]. Sequences were aligned with MUSCLE (v3.7) using default mode [79]. After alignment, ambiguous regions (i.e. containing gaps and/or poorly aligned) were removed with Gblocks (v0.91b) [80]. The phylogenetic tree was reconstructed using the maximum likelihood method implemented in the CLC Main Workbench 7. The Bishop-Friday protein substitution model was selected [81]. Reliability for internal branch was assessed using the bootstrapping method (100 bootstrap replicates).

### Antibody production

Custom polyclonal rabbit SID-2 antibody (A90) was generated by SDIX (USA) by cDNA injection coding for the intracellular domain of *C. elegans* SID-2 (S1C Fig).

### Immunostaining

Intestines of one day-old adult animals were dissected and fixed for ten minutes with 1% formaldehyde and freeze cracked on poly-lysine coated microscope slides. Dissected intestines were fixed in −30 °C methanol for 5 minutes. Fixed samples were washed with PBS-T (PBS supplemented with 1% Tween-20) prior to primary antibody addition. Primary antibodies were incubated with the samples at 4 °C for overnight. Rabbit SID-2 antibody (custom, 1:2000) was used at 1:2000 dilution, chicken GFP antibody at 1:500 (Abacm, ab13970). Secondary antibodies were incubated at 37 °C for one hour in the dark. Secondary antibodies were used anti-Rabbit-Alexa Fluor 594 (Thermofisher, A-11012, 1:1000) and anti-Chicken Alexa Fluor 488 (Jacksonimmuno, 703-545-155). Dissected and stained intestine were mounted with Vectorshield antifading agent supplemented with DAPI.

### Protein extraction

Synchronised populations of *C. elegans* were grown on 90 mm NGM agar plates and after several washes in M9 buffer and one final wash in cold 50 mM Tris pH 7.5 150 mm NaCl 0.5 mM EDTA, 0.5% NP40 and Complete Proteinase Inhibitor Cocktail (Roche) the majority of liquid was removed from the pellet and animals were snap-frozen in liquid nitrogen. To generate lysate, samples were homogenised using a Bioruptor Twin (Diagenode) and solution was cleared of debris by centrifugation at 16,000g, 4 °C for 20 minutes. Protein concentration was determined using Bradford protein assay reagent (Sigma), measuring absorbance at a wavelength of 595 nm on a UV/Visible spectrophotometer (Ultrospec 2100 pro, Amersham Biosciences) and calculating concentration of samples relative to a BSA standard curve.

### SDS-PAGE

For tris-tricine sodium dodecyl sulphate polyacrylamide gel electrophoresis (SDS-PAGE), separating gels were prepared using 330 mM Tris-HCl pH 8.45, 0.1% SDS, 8% polyacrylamide mixture (Protogel30%: 0.8% w/v acrylamide:bisacrylamide, National Diagnostics), 0.1 % ammonium persulphate (APS) (Sigma Aldrich) and 0.1 % TEMED (Sigma). Isopropanol was used to cover separating gel during polymerization and removed by washing with ddH2O. Stacking gels were poured using 330 mM Tris-HCl pH 8.45, 0.1% SDS, 5% polyacrylamide mixture, 0.1% APS and 0.1% TEMED. 50–100 µg of sample protein extract were supplemented with 2x SDS sample buffer (100 mM Tris-HCl pH 6.8, 20% glycerol, 4% SDS, 20% *β*- Mercaptoethanol, 0.2% (w/v) bromophenol blue) and denatured at 95 °C for 5 minutes. Gel electrophoresis was performed in a Min Gel Tank Life Technology system in 1x Tris Tricine buffer (T1165-500ML Sigma Aldrich Co) at 150 V samples were left to migrate until the protein ladder (PageRuler Plus pre-stained protein ladder 10–250 kDA, Fermentas) had reached the correct position.

### Western Blot

Transfer of proteins from SDS gels onto PDFM membrane (Hybond ECL, Amersham) was performed in transfer buffer (25 mM Tris-base, 190 mM glycine, 20% methanol) for 1 hour at 4 °C in a BioRad Mini Trans Blot apparatus at 250 mA. Membranes were blocked in 5% non-fat dry milk in TBS-T buffer for 60 minutes at room temperature. Primary antibody incubation was performed in a fresh batch of milk solution with antibody at appropriate dilutions over night at 4 °C with shaking. After three washes in TBST-T for 10 minutes, secondary antibody diluted in milk/TBS-T was added and membranes were incubated for one to two hours at room temperature. After 3 washes in TBS-T, bands were detected by using Immobilon Western Chemi-luminescent HRP Substrate (Millipore) according to manufacturer’s instructions, exposure of medical X-ray films (Super Rx, Fuji) to luminescent membrane and development of films on a Compact X4 automatic X-ray film processor (Xograph Imaging Systems Ltd). Primary antibodies used were: purified custom rabbit SID-2 antibody at 1:2000, purified monoclonal mouse *α*-tubulin clone DM1A (Sigma Aldrich) 1:10000. Secondary antibodies: ECL anti-mouse IgG HRP from sheep, ECL anti-rabbit IgG HRP from donkey (both GE Healthcare).

### Confocal microscopy

Images were taken on Leica SP5 confocal microscope at 63x/1.4 HCX PL Apo CS Oil. The anti-Rabbit-Alexa Fluor 594 excited using a 561nm laser and and anti-Chicken Alexa Fluor 488 was excited using a 488nm laser.

### Pyrimidine supplementation assay

IPTG-CARB NGM plates were seeded with HT115 bacteria carrying either a plasmid (pR70 Δ T7 promoter (none), L4440 (short) or GPF (long)). For the uracil condition, IPTG-CARB NGM plates were supplemented with 0.5 % uracil according to [67] and were seeded with bacteria carrying the plasmid without T7 sides. Animals were grown to adulthood and transferred for egg laying to a new plate. On the next day, 100 hundred eggs were transferred to a new plate and 24 hours later the number of hatching eggs were counted.

### Molecular cloning

Plasmid pR70 for bacteria expression ‘none’ was cloned from L4440 using Gibson cloning using home made Gibson mix using the NEB Gibson Assembly protocol (E5510) using the primer M7827 and M7828. A gene product from IDT corresponding to the SID-2 genomic sequence starting from 980 bp upstream of SID-2 start codon to 119 bp after the stop codon was cloned using Gibson cloning into using PCR linearised pUC19 with the primers M9415 and M9416. Plasmids were transformed into DH5*α* (Bioline, BIO-85026) and purified using PureLink HQ Mini Plasmid DNA (Life Technologies Ltd, K210001).

### Mutagenesis

CRISPRCas9 genome editing was performed with home made Cas9 and gRNA, crRNA from Dharmacon using the concentrations indicated in (Table. 1) according to [82]. For *C. briggsae sid-2* CRISPR dpy-10 crRNA was replaced with plasmid pCFJ90 (myo2:mcherry) and dpy-10 HR with H_2_O. *C. elegans* SX3237*sid-2(mj465)* were injection plasmid mix indicated in (Table. 2) to generate the *sid-2* rescue strain SX3432*sid-2(mj465) III; mjEx597*[*myo2::mCherry;sid-2genomic*]. The pEM2118 plasmid was omitted in the injection mix for the generation of the control strain SX3432*sid-2(mj465) III; mjEx596*[*myo2::mCherry*].

**Table 1.** CRISPR injection mix.

|  |  |
| --- | --- |
| KCl (0.5M) | 0.5 μL |
| Hepes pH 7.4 (100 mM) | 0.74 μL |
| tracrRNA (4μg/μL) | 2.5 μL |
| dpy-10 crRNA (4 μL/μL) | 0.4 μL |
| dpy-10 HR (250 ng/μL) | 0.54 μL |
| target crRNA1 (8 μg/μL) | 0.2 μL |
| target crRNA2 (8 μg/μL) | 0.2 μL |
| HR (12.5 μM) | 0.4 μL |
| Cas9 (25 μg/μL) | 0.67 μL |
| H2O | 3.85 μL |

**Table 2.** Extra-chromosomal array injection mix.

|  |  |
| --- | --- |
| pCFJ90 (myo2:mcherry) | 5 ng/μL |
| pEM2118(SID2g) | 30 ng/μL |
| 1 kb DNA Ladder | 95 ng/μL |

### RNA quantification

RNA was isolated from bacteria using TRIZOL and quantified using Qubit RNA BR Assay Kit (Life technology, Q10210). To quantify over-expressed RNA, 50 ng of total RNA was size separated using RNA ScreenTape Analysis (Agilent, 5067-5576) and quantified using TapeStation software.

### RNA library preparation

Ten L4 animals were grown for 5 days at 20 °C. After washing thoroughly with M9 to remove bacteria, eggs were isolated using bleach and resuspended in TRIsure (Bioline, BIO-38033). Embryos were lysed with 5 freeze-thaw cycles in liquid nitrogen. Total RNA was isolated by chloroform extraction. Ribosomal RNA was depleted from total RNA using NEBNext rRNA Depletion (NEB, #E6350) and libraries prepared using NEBNext Ultra™ II Directional RNA Library Prep Kit for Illumina (NEB, #E7760). Libraries were sequenced on Illumina HiSeq 1500.

### Bioinformatic analysis

RNA reads were aligned using STAR against the *C. elegans* genome WS235 or the *E. coli* HUSEC2011CHR1 genome [83]. Read counts per genetic element of the Wormbase genome annotation WS235 were calculated using feature counts [84]. Reads were normalised using pseudoreference with geometric mean row by row [85] and statistical analysis was performed using Benjamini-Hochberg (BH) adjustment [86] using the MATLAB (Mathworks) function ‘nbintest’ with the ‘VarianceLink’ setting ‘LocalRegression’. Boeck et al. embryonic RNA sequencing data was obtained from NCBI Sequence Read Archive.

### Gene ontology analysis

Significantly differentially expressed genes (FDR *<*0.01) between wild-type and *sid-2* were used as input for the enrichment Analysis of wormbase.org [68].

### Principal components analysis of embryo data

Normalised read counts from Boeck et al. embryonic, *sid-2* mutant and wild-type animal RNA sequencing data were log transformed and centred. Next, the covariance matrix and the first ten eigenvectors were calculated using the function eigs of MATLAB (Mathworks) for the Boeck et al,. embryonic RNA sequencing data alone. The projection onto the first ten eigenvectors was calculated for the centred data of all samples.

### Development and phenotype analysis

Growth curves were estimated from long-term video imaging. To obtain synchronised embryos, 20 L4 animals were transferred to a new HB101 NGM plate. After 24 hours, the now adult animals were moved to a new HB101 NGM plate and allowed to lay eggs for one hour. Next, individual eggs were transferred to imaging plates. A custom camera system was used to record back-lit images through the development from the ex utero egg stage to the egg-laying adult stage (*≈* 65 hours). To accomplish this, an imaging system was built with a robot arm mounted camera (Flea3 3.2MP monochrome, Point Grey) moving between wells to record images sequentially. Each well contained at the start a single *C. elegans* embryo nematodes and every *≈* 3 minutes a picture of the animals was recorded for *≈* 3 days. The resulting movies were analysed off-line with a custom written MATLAB script (Mathworks) to calculate a growth curve estimated by a logistic function (logistic max, logistic rate, logistic shift). Ex-utero development time was calculated using the time adults were moved to HB101 plate and the time of hatching extracted from recorded images. Similarly, time from hatching until the first egg was calculated using the time of hatching and the time of the appearance of the first laid egg extracted from recorded images. phenotypic data was analysis in a lower dimensional phenotypic space. The lower dimensional phenotypic space was constructed by calculating the covariance matrix and and the first three eigenvectors using the function ‘eigs’ of MATLAB (Mathworks). The projection of the phenotypic data was calculated for all combinations of the first three principle components. The probability density was estimated by a kernel density function with a Gaussian kernel.

### Imaging plates

Imaging plates for developmental analysis were made using the standard NGM recipe, but without peptone and cholesterol. Furthermore, agarose was substituted with 0.8% Gelzan (Sigma G1910-250G) for a more transparent gel. Imaging plates were seeded with with one *µ*L of HB101 concentrate at optical density 20.

### L1 length analysis

To obtain synchronised embryos for L1 length analysis, 40 L4 animals were transferred to a new HB101 NGM plate. After 24 hours, the now adult animals were moved to a new HB101 NGM plate and allowed to lay eggs for one hour. 50 eggs were transferred on imaging plates without food and imaged with the the custom imaging system. The resulting movies were analysed off-line with a custom written MATLAB script (Mathworks). In short, animals were segmented from the background using an intensity threshold and a skeleton was extracted using the *bwmoprh* function. The length of the skeleton was computed using the *bwdistgeodesic* function.

## Supporting information

**S1.**
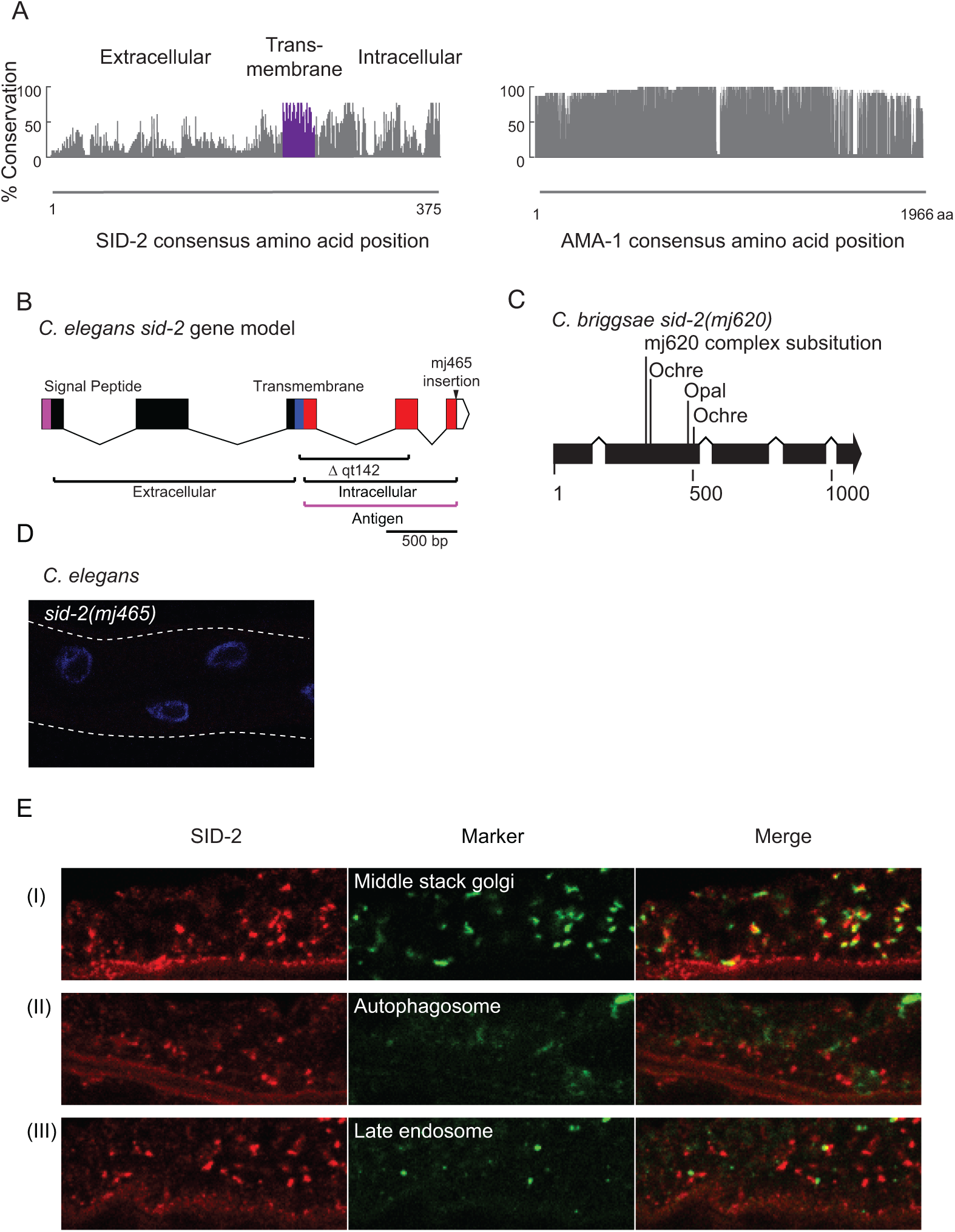
Conservation analysis of SID-2, gene model of *sid-2* and SID-2 localises next to the middle Golgi stack. A) Plot showing the conservation along the protein sequence for a alignment of 25 SID-2 protein sequences (left) and 23 RNA polymerase II subunit A protein sequences (right). B) Gene model of *C. elegans’ sid-2*. Deleted region in the allele *sid-2(qt142)* and insertion in *sid-2(mj465)* (triangle) are indicated. In addition, coding regions for protein domains (Signal peptide (purple), extra-cellular (green), trans-membrane (blue) and intra-cellular domain (red)) are highlighted. Predicted SID-2 antibody target side indicated by a bar (magenta) corresponds to the intra-cellular domain. C) Gene model of *C. briggsae’s sid-2*. Complex substitution of the allele *sid-2(mj600)* causes a frame shift. The three closest premature STOP codons to the complex substitution are indicated. D) Confocal microscopy image of *C. elegans sid-2(mj465)* mutant dissected adult intestines after SID-2 immunofluorescent staining. DNA was stained using DAPI (blue). SID-2 was immunostained using SID-2 antibody. E) Confocal microscopy image of *C. elegans* dissected adult intestines expressing GFP fused to cellular compartment markers after SID-2 and GFP co-immunofluorescent staining. DNA was stained using DAPI (blue). SID-2 and GFP were immunostained using SID-2 antibody and GFP antibody, respectively. Left column shows SID-2 antibody staining in red. Middle column show GFP staining in green. Right column shows the SID-2 and GFP merged image. (I) Middle stack Golgi (MANS::GFP), (II) autophagosome (LGG-1::GFP) and (III) late endosome (GFP::RAB-7), representative image is shown (n = 3).

**S2.**
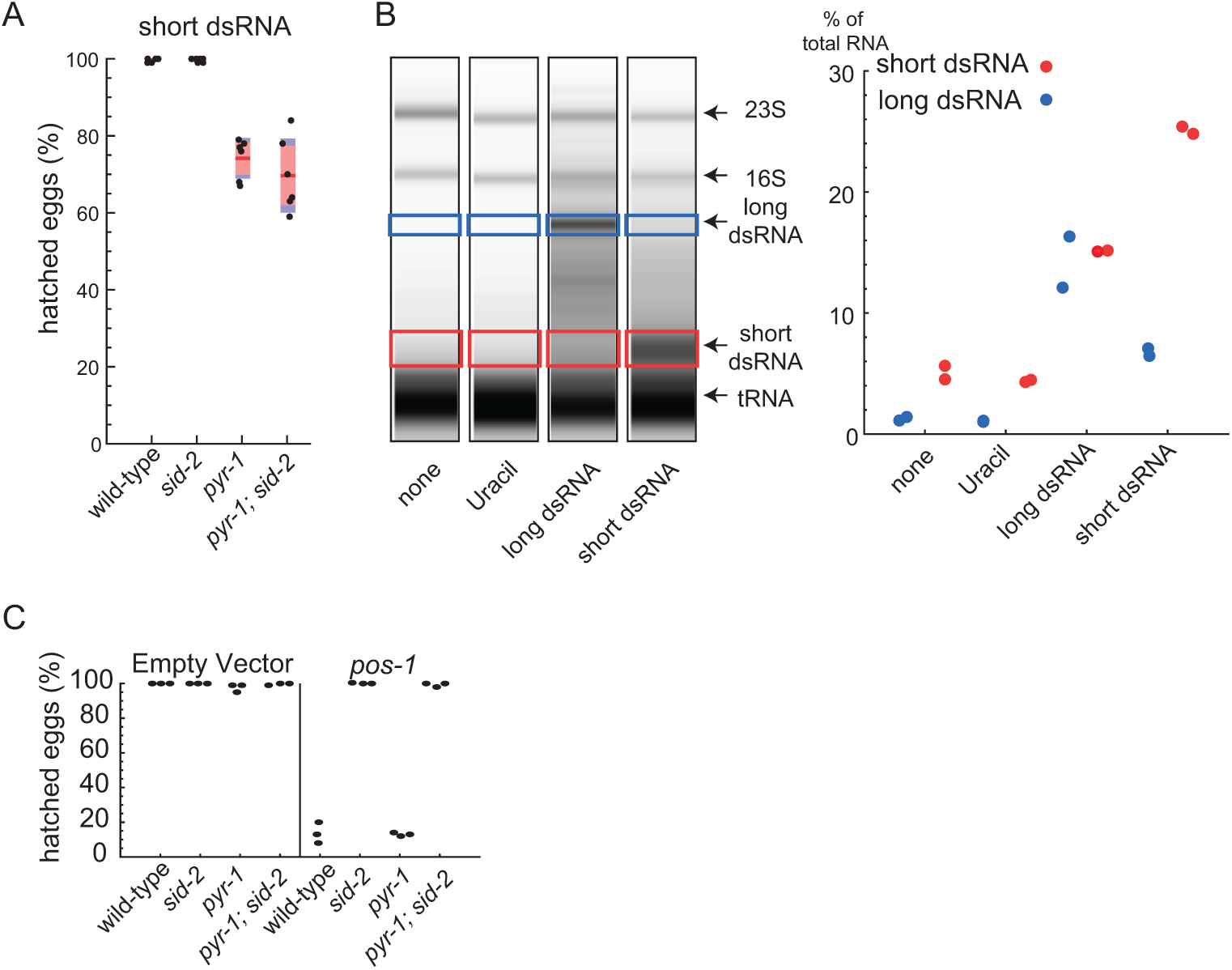
Long and short dsRNA are present at high levels in the environment and *pyr-1* mutants are dsRNA uptake competent. A) Embryonic hatching after feeding short dsRNA. Black dots represent hatching efficiency of eggs laid by adult animals. Three biological experiments with 2 technical replicates for each genotype was performed. Red lines indicate the mean. Red boxes indicate the 95% confidence interval of the mean. Blue boxes the standard deviation. B) RNA electrophoresis of bacteria total RNA using TapeStation. Total RNA was isolated from bacteria used in RNA feeding assays and equal amounts of total RNA were assayed. Quantity of RNA in blue (long dsRNA) and red (short dsRNA) indicated area were plotted. Data for two biological replicates is shown. Red lines indicate the mean. Red boxes indicate the 95% confidence interval of the mean. C) *Pos-1* RNAi by feeding assay in the presence of uracil. The ability of dsRNA uptake of *pyr-1* mutant animals was accessed in an RNAi by feeding assay. Eggs of wild-type, *sid-2*, *pyr-1*, or *pyr-1;sid-2* double mutant animals were examined after the parents were raised from L1 to adult stage on bacteria expressing negative control dsRNA or *pos-1* dsRNA. Black dots represent hatching efficiency of eggs from adult animals. Three biological replicates for each genotype were performed.

**S3.**
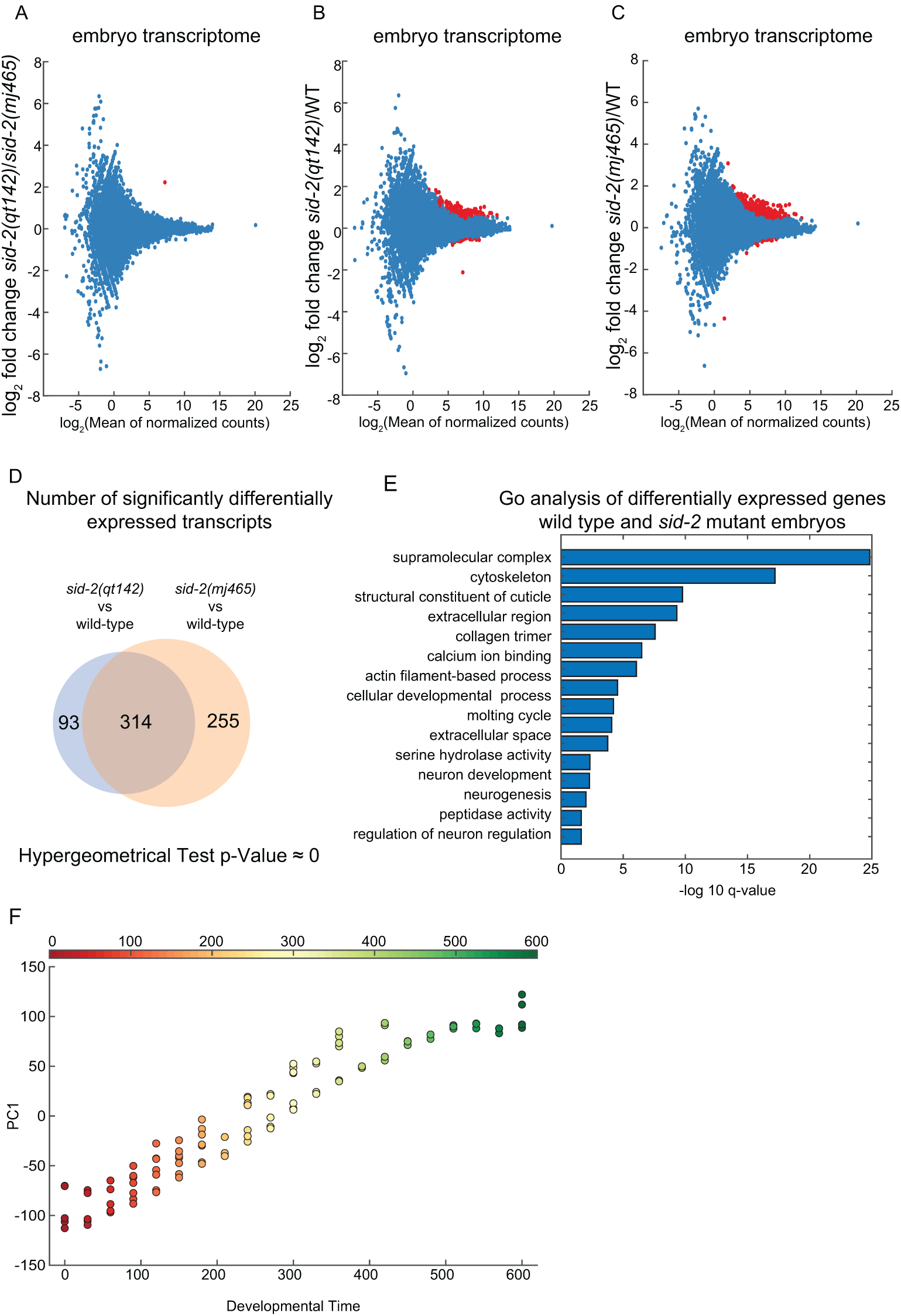
Differentially expressed genes of *sid-2* mutant embryos are associated with developmental gene ontologies. A) MA plot comparing the transcriptome of *sid-2(qt142) (n = 5)* and *sid-2(mj465)* (n = 4) mutant embryos. Each red circle represents a statistically significant DE transcripts (FDR *<*0.01). B) MA plot comparing the transcriptomes of wild type (n = 5 and *sid-2(qt142)* (n = 5) mutant embryos. Each red circle represents a statistically significant DE transcript (FDR *<*0.01). C) MA plot comparing the transcriptome of wild type (n = 5) and *sid-2(mj465)* (n = 4) mutant embryos. Each red circle represents a statistically significant DE transcript (FDR *<*0.01). D) Venn diagram showing the overlap of the transcripts of *sid-2(qt142)* and *sid-2(mj465)* embryos compared to wild type. Hyper-geometric Test for the overlap (p *≈* 0). E) Gene ontology analysis of DE transcripts between wild type and *sid-2* mutant embryos. F) Scatter plot correlating developmental time and position along the first principal component for individual embryo transcriptome samples.

**S4.**
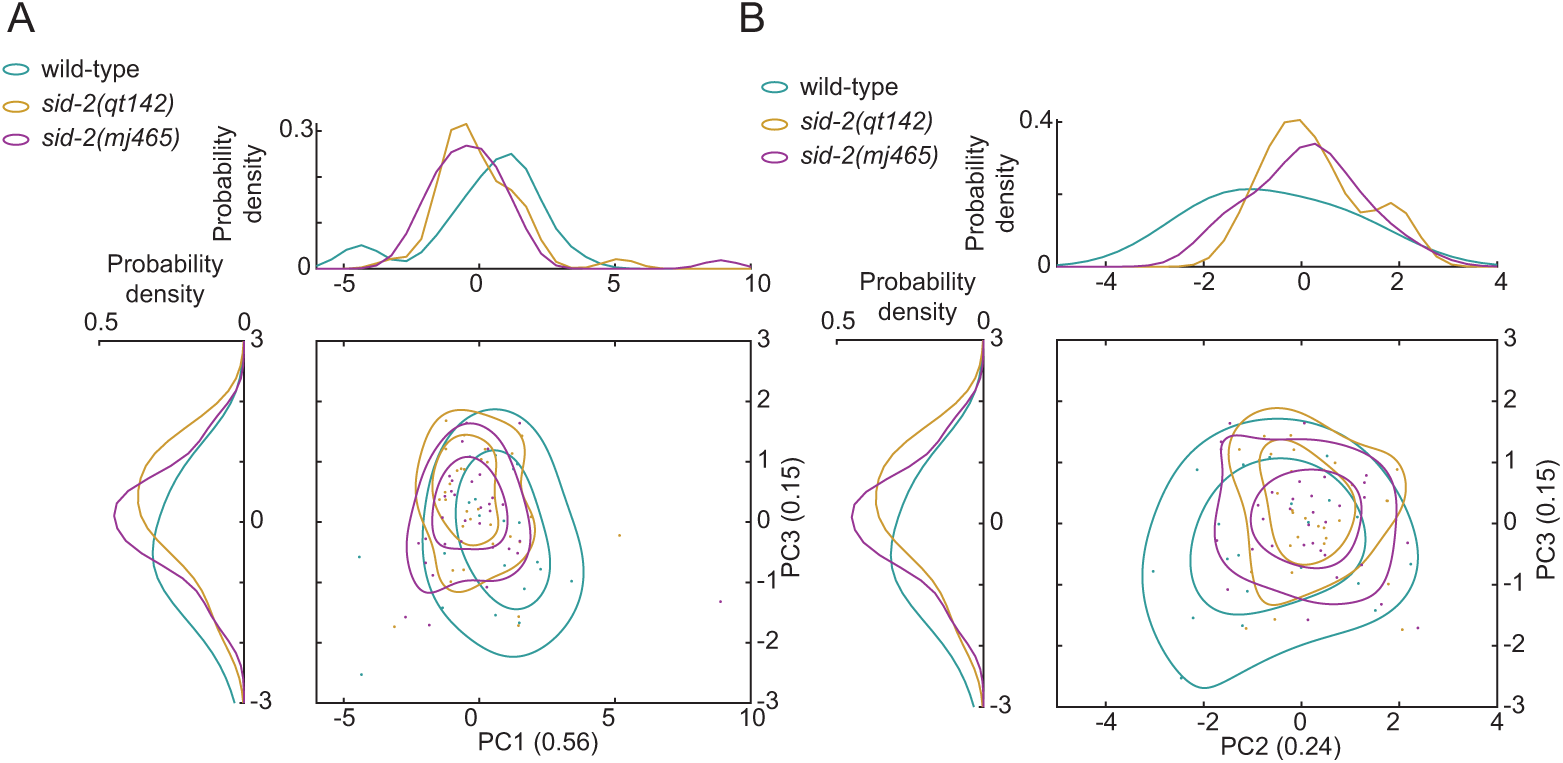
Principal components analysis representing the combined phenotypic. A) Principal components analysis representing the combined phenotypic data along the first and third and B) the second and third principal component of the analysed wild-type animals (n = 29) and *sid-2(qt142)* (n = 30), *sid-2(mj465)* (n = 31) mutants. Individual circles represent aggregated phenotypic data of a individual animal. The line represent the 33% and 66% contour line. Probability density estimate of the phenotypic data are plotted to the left and above of the PCA plot.

**S5.**
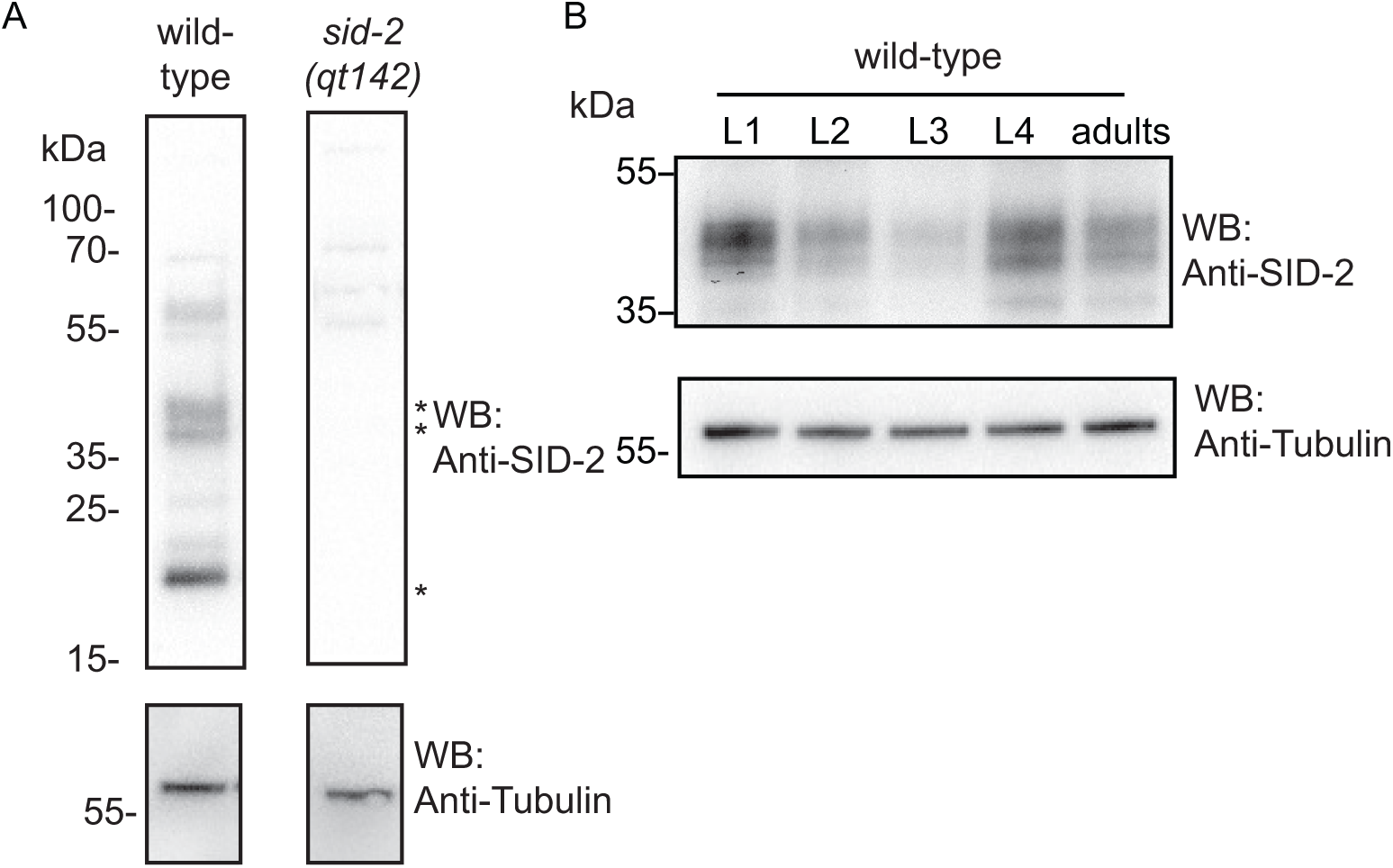
SID-2 is expressed in all larvae stages and adults. A) Specificity of SID-2 antibody was tested by western blot analysis of wild-type and *sid-2(qt142)* animal lysate (representative blot is shown (n = 2), Full blots are shown in S8 Fig. B) SID-2 expression through development. Top: SID-2 western blot of wild-type animal lysate of different developmental stages using SID-2 antibody (SID-2 isoforms are indicated with a *). Bottom: Tubulin immunoblotting of the above membrane (representative blot is shown (n = 2). Full blots are shown in S8 Fig.

**S6.**
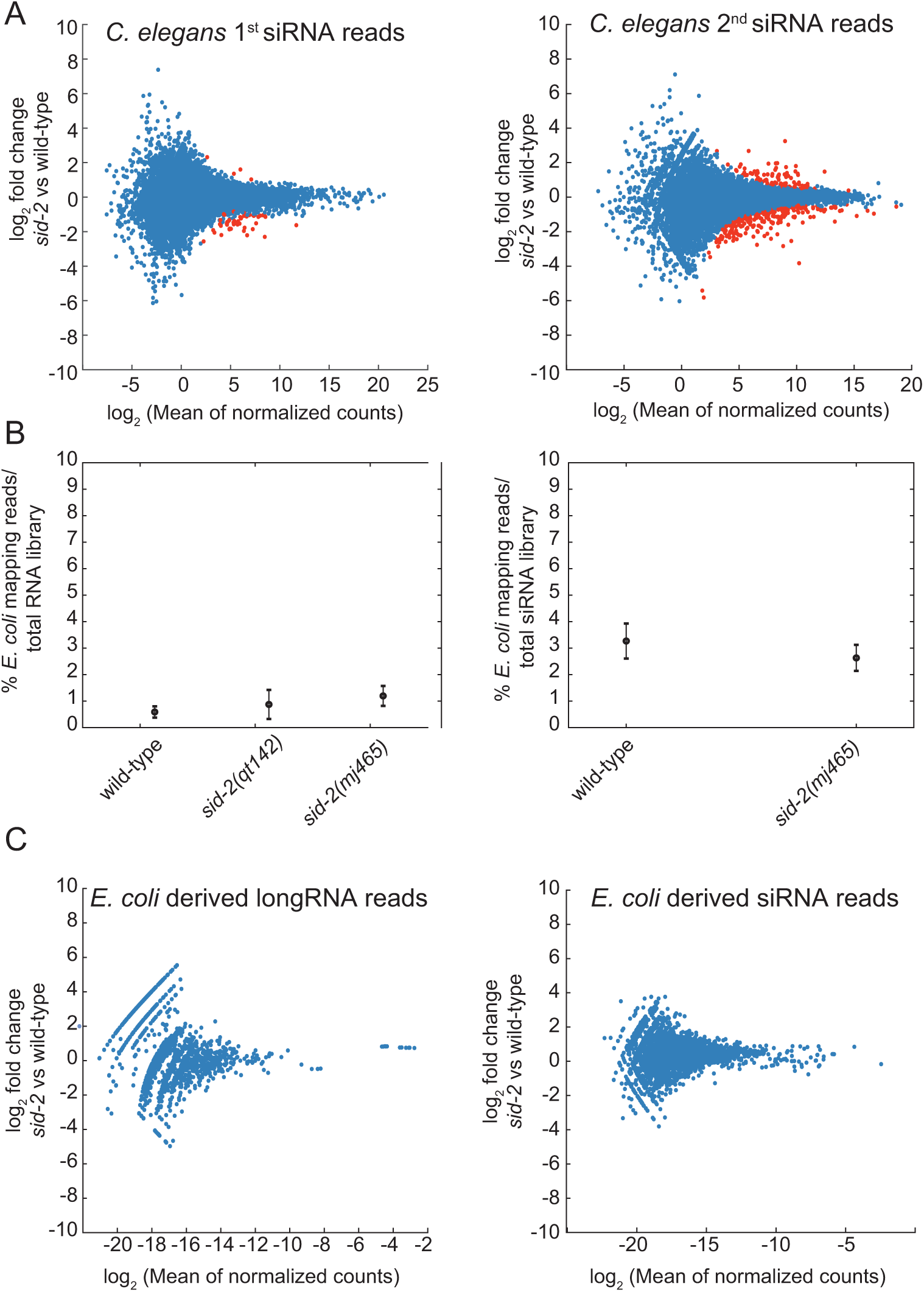
No differentially abundant *E. coli* RNA were detected by RNA sequencing. A) MA plot visualising (left) primary siRNAs of *C. elegans* L4 wild-type animals (n = 4) and *sid-2* mutants (*sid-2(mj465)* n = 5) and (right) secondary siRNAs of wild type (n = 3) and *sid-2* mutants (total n = 6, *sid-2(qt142)* n = 3, *sid-2(mj465)* n = 3). Each red circle represents a statistically significant DE transcript (FDR *<*0.01). B) Plot visualising (left) the percentage of *E. coli* derived RNA reads in wild-type (n = 3) and indicated *sid-2* mutants (total n = 6 *sid-2(qt142)* n = 3, *sid-2(mj465)* n = 3) and (right) the percentage of *E. coli* derived siRNA reads in wild-type (n = 4) and indicated *sid-2(mj465)* mutants (n = 5). Bars indicate standard deviation of the mean, dots indicate the mean. C) MA plot visualising (left) *E. coli* long RNA associated with *C. elegans* wild-type animals (n = 3) and *sid-2* mutants (total n = 6, *sid-2(qt142)* n = 3, *sid-2(mj465)* n = 3) and (right) *E. coli* short RNAs associated with L4 wild type (n = 4) and *sid-2* mutants (*sid-2(mj465)* n = 5). Each red circle represents a statistically significant DE transcript (FDR *<*0.01).

**S7.**
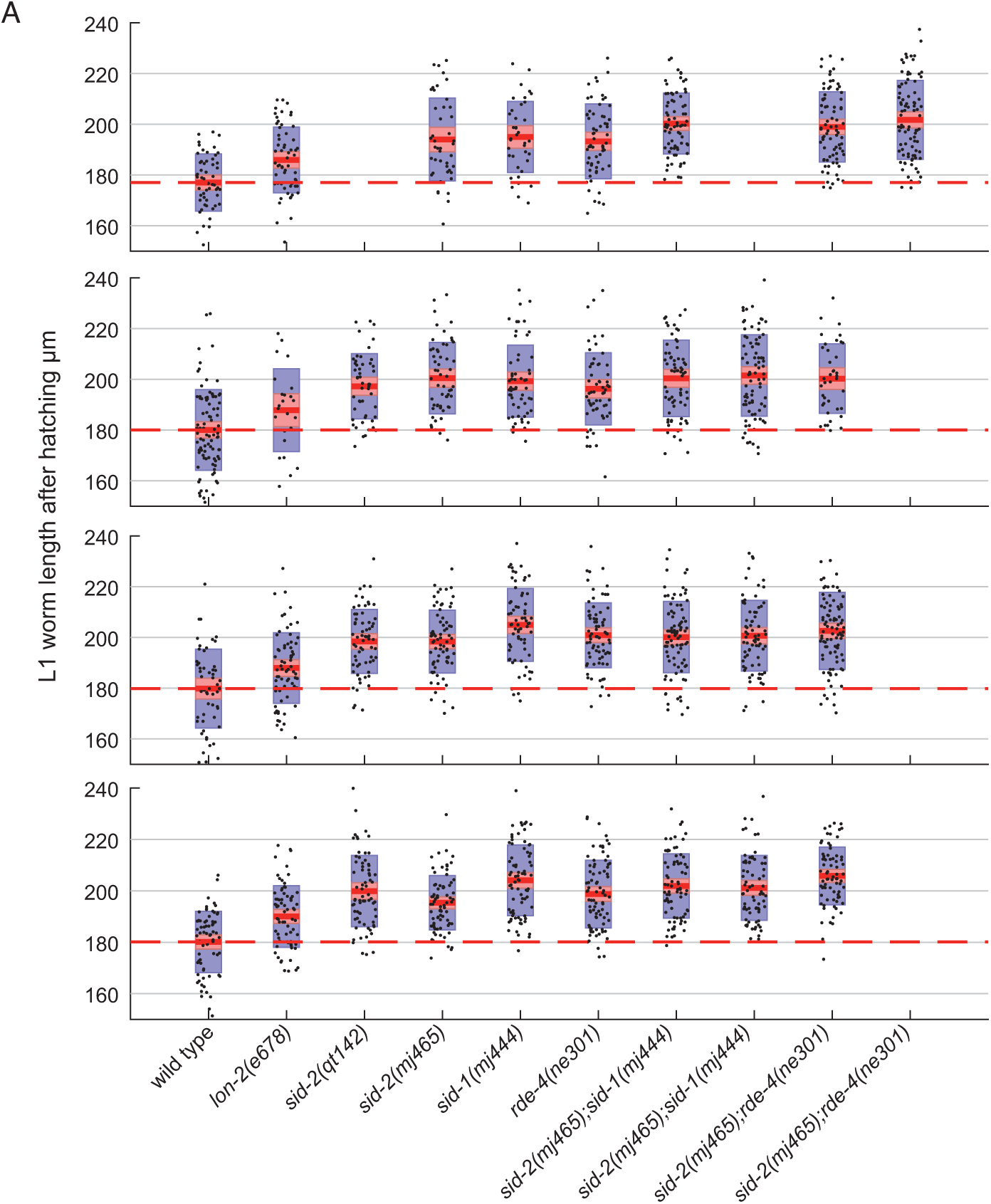
Individual experiments measuring L1 larva length at hatching. A) body length at hatching of *C. elegans* wild-type and indicated mutant animals. Black dots represent body length of one hatched animal. Red lines indicate the mean. Red boxes indicate the 95% confidence interval of the mean. Blue boxes indicate the standard deviation.

**S8.**
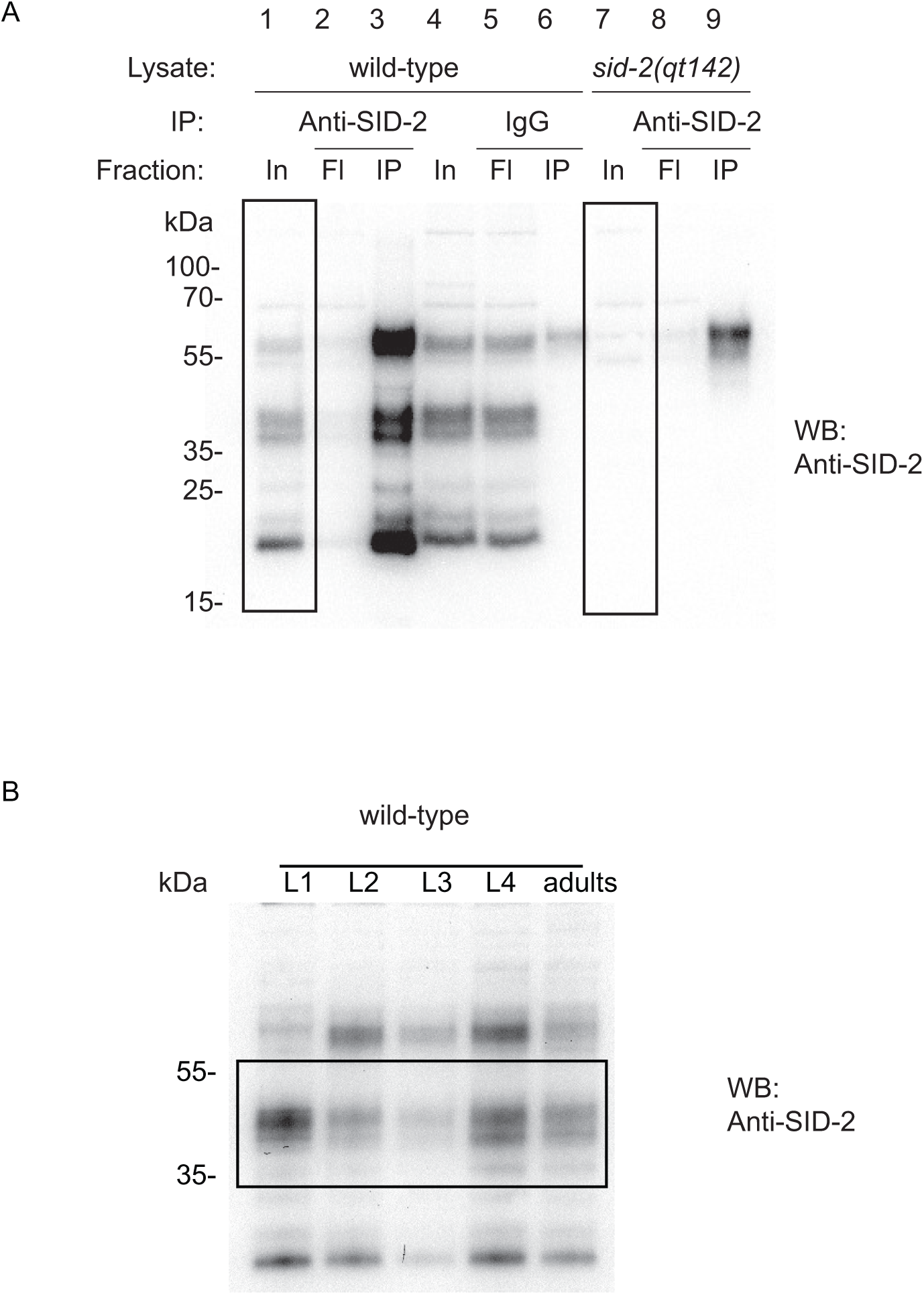
Images of full western blot membrane. A) Full western blot for S5A Fig. B) Full western blot for S5B Fig

**S1 Table. SID-2 and AMA-1 protein sequences in *Caenorhabditis*.**

**S2 Table. *sid-1* and *sid-2* animals are resistant to *rpb-2* RNAi by feeding.** RNAi feeding experiment on L4 larva fed with either control (L4440) or *rpb-2* RNAi. The presents (+) or absence (−) of F1 larvae was scored after 48. Three independent experiments were conducted.

**S3 Table. Phenotypic data and variance analysis**

**S4 Table. Strains used in this study**

**S5 Table. RNAi clones used in this study**

**S6 Table. Primer used in this study**

**S7 Table. Plasmids used in this study**

**S1 Data. SID-2 and AMA-1 alignment**

**S2 Data. Allele maps**

## Acknowledgements

We are grateful to Craig Hunter, Julie Ahringer, Chiara Cerrato and Yan Dong for sharing strains prior to publication. Alper Akay for sharing a plasmid. Nicolas Bologna, Giulia Furlan, Alexandra Bezler, Melanie Tanguy, Alexandra Dellaire, Navin B Ramakrishna and the anonymous reviewers for comments on the manuscript. The Miska lab members and Kin Man Suen for advice. We thank Kay Harnish of the Gurdon Institute Sequencing Facility for managing the high-throughput sequencing. The Julie Ahringer laboratory and the Miska laboratory members for comments on the manuscript and advice. Juanita Baker Hay for media preparation and Marc Ridyard for maintenance of the worm collection. Some strains were provided by the *Caenorhabditis* Genetics Center (CGC) and the National BioResource Project (NBRP).

## Availability of data

Sequencing data is available in the European Nucleotide Archive under the study accession number PRJEB32813.

## Author’s Contributions

Fabian Braukmann, David Jordan contributed to experimental design, collected data, analysed and interpreted results, and drafted the manuscript. Eric Alexander Miska contributed to design of experiments and analyses, results interpretation and drafting the manuscript. All authors read and approved the final manuscript.

## References

1. Knip M, Constantin ME, Thordal-Christensen H. Trans-kingdom cross-talk: small RNAs on the move. PLoS genetics. 2014;10(9):e1004602.

2. Zhou G, Zhou Y, Chen X. New insight into inter-kingdom communication: horizontal transfer of mobile small RNAs. Frontiers in microbiology. 2017;8:768.

3. Timmons L, Fire A. Specific interference by ingested dsRNA. Nature. 1998;395(6705):854.

4. Timmons L, Court DL, Fire A. Ingestion of bacterially expressed dsRNAs can produce specific and potent genetic interference in Caenorhabditis elegans. Gene. 2001;263(1-2):103–112.

5. Nowara D, Gay A, Lacomme C, Shaw J, Ridout C, Douchkov D, et al. HIGS: host-induced gene silencing in the obligate biotrophic fungal pathogen Blumeria graminis. The Plant Cell. 2010;22(9):3130–3141.

6. Koch A, Kumar N, Weber L, Keller H, Imani J, Kogel KH. Host-induced gene silencing of cytochrome P450 lanosterol C14*α*-demethylase–encoding genes confers strong resistance to Fusarium species. Proceedings of the National Academy of Sciences. 2013;110(48):19324–19329.

7. Baum JA, Bogaert T, Clinton W, Heck GR, Feldmann P, Ilagan O, et al. Control of coleopteran insect pests through RNA interference. Nature biotechnology. 2007;25(11):1322.

8. Mulot M, Boissinot S, Monsion B, Rastegar M, Clavijo G, Halter D, et al. Comparative analysis of RNAi-based methods to down-regulate expression of two genes expressed at different levels in Myzus persicae. Viruses. 2016;8(11):316.

9. Wang M, Weiberg A, Lin FM, Thomma BP, Huang HD, Jin H. Bidirectional cross-kingdom RNAi and fungal uptake of external RNAs confer plant protection. Nature plants. 2016;2(10):16151.

10. Koch A, Biedenkopf D, Furch A, Weber L, Rossbach O, Abdellatef E, et al. An RNAi-based control of Fusarium graminearum infections through spraying of long dsRNAs involves a plant passage and is controlled by the fungal silencing machinery. PLoS pathogens. 2016;12(10):e1005901.

11. Buck AH, Coakley G, Simbari F, McSorley HJ, Quintana JF, Le Bihan T, et al. Exosomes secreted by nematode parasites transfer small RNAs to mammalian cells and modulate innate immunity. Nature communications. 2014;5:5488.

12. Weiberg A, Wang M, Lin FM, Zhao H, Zhang Z, Kaloshian I, et al. Fungal small RNAs suppress plant immunity by hijacking host RNA interference pathways. Science. 2013;342(6154):118–123.

13. Zhu K, Liu M, Fu Z, Zhou Z, Kong Y, Liang H, et al. Plant microRNAs in larval food regulate honeybee caste development. PLoS genetics. 2017;13(8):e1006946.

14. Kamath RS, Ahringer J. Genome-wide RNAi screening in Caenorhabditis elegans. Methods. 2003;30(4):313–321.

15. Zotti M, dos Santos EA, Cagliari D, Christiaens O, Taning CNT, Smagghe G. RNA interference technology in crop protection against arthropod pests, pathogens and nematodes. Pest management science. 2018;74(6):1239–1250.

16. Mitter N, Worrall EA, Robinson KE, Li P, Jain RG, Taochy C, et al. Clay nanosheets for topical delivery of RNAi for sustained protection against plant viruses. Nature plants. 2017;3(2):16207.

17. Hunter W, Ellis J, Hayes J, Westervelt D, Glick E, Williams M, et al. Large-scale field application of RNAi technology reducing Israeli acute paralysis virus disease in honey bees (Apis mellifera, Hymenoptera: Apidae). PLoS pathogens. 2010;6(12):e1001160.

18. Ramesh S. Non-coding RNAs in crop genetic modification: considerations and predictable environmental risk assessments (ERA). Molecular biotechnology. 2013;55(1):87–100.

19. Bachman PM, Bolognesi R, Moar WJ, Mueller GM, Paradise MS, Ramaseshadri P, et al. Characterization of the spectrum of insecticidal activity of a double-stranded RNA with targeted activity against Western Corn Rootworm (Diabrotica virgifera virgifera LeConte). Transgenic research. 2013;22(6):1207–1222.

20. Auer C, Frederick R. Crop improvement using small RNAs: applications and predictive ecological risk assessments. Trends in biotechnology. 2009;27(11):644–651.

21. Ramon M, Devos Y, Lanzoni A, Liu Y, Gomes A, Gennaro A, et al. RNAi-based GM plants: food for thought for risk assessors. Plant biotechnology journal. 2014;12(9):1271–1273.

22. Casacuberta JM, Devos Y, Du Jardin P, Ramon M, Vaucheret H, Nogue F. Biotechnological uses of RNAi in plants: risk assessment considerations. Trends in Biotechnology. 2015;33(3):145–147.

23. Roberts AF, Devos Y, Lemgo GN, Zhou X. Biosafety research for non-target organism risk assessment of RNAi-based GE plants. Frontiers in plant science. 2015;6:958.

24. Khajuria C, Ivashuta S, Wiggins E, Flagel L, Moar W, Pleau M, et al. Development and characterization of the first dsRNA-resistant insect population from western corn rootworm, Diabrotica virgifera virgifera LeConte. PloS one. 2018;13(5):e0197059.

25. Whangbo JS, Hunter CP. Environmental RNA interference. Trends in genetics. 2008;24(6):297–305.

26. Winston WM, Sutherlin M, Wright AJ, Feinberg EH, Hunter CP. Caenorhabditis elegans SID-2 is required for environmental RNA interference. Proceedings of the National Academy of Sciences. 2007;104(25):10565–10570.

27. McEwan DL, Weisman AS, Hunter CP. Uptake of extracellular double-stranded RNA by SID-2. Molecular cell. 2012;47(5):746–754.

28. Saleh MC, van Rij RP, Hekele A, Gillis A, Foley E, O’Farrell PH, et al. The endocytic pathway mediates cell entry of dsRNA to induce RNAi silencing. Nature cell biology. 2006;8(8):793.

29. Winston WM, Molodowitch C, Hunter CP. Systemic RNAi in C. elegans requires the putative transmembrane protein SID-1. Science. 2002;295(5564):2456–2459.

30. Feinberg EH, Hunter CP. Transport of dsRNA into cells by the transmembrane protein SID-1. Science. 2003;301(5639):1545–1547.

31. Wang E, Hunter CP. SID-1 Functions in Multiple Roles To Support Parental RNAi in Caenorhabditis elegans. Genetics. 2017;207(2):547–557.

32. Tabara H, Yigit E, Siomi H, Mello CC. The dsRNA binding protein RDE-4 interacts with RDE-1, DCR-1, and a DExH-box helicase to direct RNAi in C. elegans. Cell. 2002;109(7):861–871.

33. Hammond SM, Bernstein E, Beach D, Hannon GJ. An RNA-directed nuclease mediates post-transcriptional gene silencing in Drosophila cells. Nature. 2000;404(6775):293.

34. Steiner FA, Okihara KL, Hoogstrate SW, Sijen T, Ketting RF. RDE-1 slicer activity is required only for passenger-strand cleavage during RNAi in Caenorhabditis elegans. Nature structural & molecular biology. 2009;16(2):207.

35. Doench JG, Petersen CP, Sharp PA. siRNAs can function as miRNAs. Genes & development. 2003;17(4):438–442.

36. Martinez J, Patkaniowska A, Urlaub H, Lührmann R, Tuschl T. Single-stranded antisense siRNAs guide target RNA cleavage in RNAi. Cell. 2002;110(5):563–574.

37. Martinez J, Tuschl T. RISC is a 5*^1^* phosphomonoester-producing RNA endonuclease. Genes & development. 2004;18(9):975–980.

38. Béthune J, Artus-Revel CG, Filipowicz W. Kinetic analysis reveals successive steps leading to miRNA-mediated silencing in mammalian cells. EMBO reports. 2012;13(8):716–723.

39. Djuranovic S, Nahvi A, Green R. miRNA-mediated gene silencing by translational repression followed by mRNA deadenylation and decay. Science. 2012;336(6078):237–240.

40. Eichhorn SW, Guo H, McGeary SE, Rodriguez-Mias RA, Shin C, Baek D, et al. mRNA destabilization is the dominant effect of mammalian microRNAs by the time substantial repression ensues. Molecular cell. 2014;56(1):104–115.

41. Jonas S, Izaurralde E. Towards a molecular understanding of microRNA-mediated gene silencing. Nature Reviews Genetics. 2015;16(7):421.

42. Sijen T, Fleenor J, Simmer F, Thijssen KL, Parrish S, Timmons L, et al. On the role of RNA amplification in dsRNA-triggered gene silencing. Cell. 2001;107(4):465–476.

43. Pak J, Fire A. Distinct populations of primary and secondary effectors during RNAi in C. elegans. Science. 2007;315(5809):241–244.

44. Sijen T, Steiner FA, Thijssen KL, Plasterk RH. Secondary siRNAs result from unprimed RNA synthesis and form a distinct class. Science. 2007;315(5809):244–247.

45. Vastenhouw NL, Brunschwig K, Okihara KL, Müller F, Tijsterman M, Plasterk RH. Gene expression: long-term gene silencing by RNAi. Nature. 2006;442(7105):882.

46. Alcazar RM, Lin R, Fire AZ. Transmission dynamics of heritable silencing induced by double-stranded RNA in Caenorhabditis elegans. Genetics. 2008;180(3):1275–1288.

47. Burton NO, Burkhart KB, Kennedy S. Nuclear RNAi maintains heritable gene silencing in Caenorhabditis elegans. Proceedings of the National Academy of Sciences. 2011;108(49):19683–19688.

48. Buckley BA, Burkhart KB, Gu SG, Spracklin G, Kershner A, Fritz H, et al. A nuclear Argonaute promotes multigenerational epigenetic inheritance and germline immortality. Nature. 2012;489(7416):447.

49. Payne S. Chapter 10 - Introduction to RNA Viruses. In: Payne S, editor. Viruses. Academic Press; 2017. p. 97–105.

50. Reich DP, Tyc KM, Bass BL. C. elegans ADARs antagonize silencing of cellular dsRNAs by the antiviral RNAi pathway. Genes & development. 2018;32(3-4):271–282.

51. Lybecker M, Zimmermann B, Bilusic I, Tukhtubaeva N, Schroeder R. The double-stranded transcriptome of Escherichia coli. Proceedings of the National Academy of Sciences. 2014;111(8):3134–3139.

52. Lehner B, Williams G, Campbell RD, Sanderson CM. Antisense transcripts in the human genome. Trends in Genetics. 2002;18(2):63–65.

53. Shendure J, Church GM. Computational discovery of sense-antisense transcription in the human and mouse genomes. Genome biology. 2002;3(9):research0044–1.

54. Lioliou E, Sharma CM, Caldelari I, Helfer AC, Fechter P, Vandenesch F, et al. Global regulatory functions of the Staphylococcus aureus endoribonuclease III in gene expression. PLoS genetics. 2012;8(6):e1002782.

55. Zinad HS, Natasya I, Werner A. Natural antisense transcripts at the interface between host genome and mobile genetic elements. Frontiers in microbiology. 2017;8:2292.

56. Culley AI, Lang AS, Suttle CA. Metagenomic analysis of coastal RNA virus communities. Science. 2006;312(5781):1795–1798.

57. Djikeng A, Kuzmickas R, Anderson NG, Spiro DJ. Metagenomic analysis of RNA viruses in a fresh water lake. PLOS one. 2009;4(9):e7264.

58. Cantalupo PG, Calgua B, Zhao G, Hundesa A, Wier AD, Katz JP, et al. Raw sewage harbors diverse viral populations. MBio. 2011;2(5):e00180–11.

59. Steward GF, Culley AI, Mueller JA, Wood-Charlson EM, Belcaid M, Poisson G. Are we missing half of the viruses in the ocean? The ISME journal. 2013;7(3):672.

60. Decker CJ, Parker R. Analysis of double-stranded RNA from microbial communities identifies double-stranded RNA virus-like elements. Cell reports. 2014;7(3):898–906.

61. Maori E, Garbian Y, Kunik V, Mozes-Koch R, Malka O, Kalev H, et al. A transmissible RNA pathway in honey bees. Cell reports. 2019;27(7):1949–1959.

62. Maori E, Navarro IC, Boncristiani H, Seilly DJ, Rudolph KLM, Sapetschnig A, et al. A Secreted RNA Binding Protein Forms RNA-Stabilizing Granules in the Honeybee Royal Jelly. Molecular cell. 2019;.

63. Nuez I, Félix MA. Evolution of susceptibility to ingested double-stranded RNAs in Caenorhabditis nematodes. PloS one. 2012;7(1):e29811.

64. Slos D, Sudhaus W, Stevens L, Bert W, Blaxter M. Caenorhabditis monodelphis sp. n.: defining the stem morphology and genomics of the genus Caenorhabditis. BMC Zoology. 2017;2(1):4.

65. Stevens L, Félix MA, Beltran T, Braendle C, Caurcel C, Fausett S, et al. Comparative genomics of ten new Caenorhabditis species. bioRxiv. 2018; p. 398446.

66. Imae R, Dejima K, Kage-Nakadai E, Arai H, Mitani S. Endomembrane-associated RSD-3 is important for RNAi induced by extracellular silencing RNA in both somatic and germ cells of Caenorhabditis elegans. Scientific reports. 2016;6:28198.

67. Franks DM, Izumikawa T, Kitagawa H, Sugahara K, Okkema PG. C. elegans pharyngeal morphogenesis requires both de novo synthesis of pyrimidines and synthesis of heparan sulfate proteoglycans. Developmental biology. 2006;296(2):409–420.

68. Angeles-Albores D, Lee RY, Chan J, Sternberg PW. Two new functions in the WormBase Enrichment Suite. microPublication Biology. 2018;.

69. Boeck ME, Huynh C, Gevirtzman L, Thompson OA, Wang G, Kasper DM, et al. The time-resolved transcriptome of C. elegans. Genome research. 2016;26(10):1441–1450.

70. Hirschberg CB, Robbins PW, Abeijon C. Transporters of nucleotide sugars, ATP, and nucleotide sulfate in the endoplasmic reticulum and Golgi apparatus; 1998.

71. Brenner S. The genetics of Caenorhabditis elegans. Genetics. 1974;77(1):71–94.

72. Bergman A, Siegal ML. Evolutionary capacitance as a general feature of complex gene networks. Nature. 2003;424(6948):549.

73. Raj A, Rifkin SA, Andersen E, Van Oudenaarden A. Variability in gene expression underlies incomplete penetrance. Nature. 2010;463(7283):913.

74. Shih JD, Fitzgerald MC, Sutherlin M, Hunter CP. The SID-1 double-stranded RNA transporter is not selective for dsRNA length. Rna. 2009;15(3):384–390.

75. Shih JD, Hunter CP. SID-1 is a dsRNA-selective dsRNA-gated channel. Rna. 2011;17(6):1057–1065.

76. Wang J, Silva M, Haas LA, Morsci NS, Nguyen KC, Hall DH, et al. C. elegans ciliated sensory neurons release extracellular vesicles that function in animal communication. Current Biology. 2014;24(5):519–525.

77. Wang J, Barr MM. Cell–cell communication via ciliary extracellular vesicles: clues from model systems. Essays in biochemistry. 2018;62(2):205–213.

78. Braukmann F, Jordan D, Miska E. Artificial and natural RNA interactions between bacteria and C. elegans. RNA biology. 2017;14(4):415–420.

79. Edgar RC. MUSCLE: multiple sequence alignment with high accuracy and high throughput. Nucleic acids research. 2004;32(5):1792–1797.

80. Castresana J. Selection of conserved blocks from multiple alignments for their use in phylogenetic analysis. Molecular biology and evolution. 2000;17(4):540–552.

81. Bishop M, Friday AE. Evolutionary trees from nucleic acid and protein sequences. Proc R Soc Lond B. 1985;226(1244):271–302.

82. Paix A, Folkmann A, Seydoux G. Precision genome editing using CRISPR-Cas9 and linear repair templates in C. elegans. Methods. 2017;121:86–93.

83. Dobin A, Davis CA, Schlesinger F, Drenkow J, Zaleski C, Jha S, et al. STAR: ultrafast universal RNA-seq aligner. Bioinformatics. 2013;29(1):15–21.

84. Liao Y, Smyth GK, Shi W. featureCounts: an efficient general purpose program for assigning sequence reads to genomic features. Bioinformatics. 2013;30(7):923–930.

85. Anders S, Huber W. Differential expression analysis for sequence count data. Genome biology. 2010;11(10):R106.

86. Author T, Benjamini Y, Hochberg Y, Benjaminit Y. Controlling the False Discovery Rate: A Practical and Powerful Approach to Multiple Controlling the False Discovery Rate: a Practical and Powerful Approach to Multiple Testing. Source JR Stat Soc Ser BJR Stat Soc Ser BMethodological) JR Stat Soc B. 1995;57:289–300.

